# Expression profiling reveals novel role of Hunchback in retinal glia cell development and blood-brain barrier integrity

**DOI:** 10.1101/114363

**Authors:** Montserrat Torres-Oliva, Julia Schneider, Gordon Wiegleb, Felix Kaufholz, Nico Posnien

## Abstract

The development of different cell types must be tightly coordinated in different organs. The developing head of *Drosophila melanogaster* represents an excellent model to study the molecular mechanisms underlying this coordination because the eye-antennal imaginal discs contain the organ anlagen of nearly all adult head structures, such as the compound eyes or the antennae. We studied the genome wide gene expression dynamics during eye-antennal disc development in *D. melanogaster* to identify new central regulators of the underlying gene regulatory network. Expression based gene clustering and transcription factor motif enrichment analyses revealed a central regulatory role of the transcription factor Hunchback (Hb). We confirmed that *hb* is expressed in two polyploid retinal subperineurial glia cells (carpet cells). Our functional analysis shows that Hb is necessary for carpet cell development and loss of Hb function results in abnormal glia cell migration and photoreceptor axon guidance patterns. Additionally, we show for the first time that the carpet cells are an integral part of the blood-brain barrier.

## Introduction

The development of complex organs is often accompanied by extensive cell- and tissue rearrangements. In some extreme cases, initially simple cells undergo profound morphological changes such as extensive cell fusions of muscle precursor cells to form syncytial muscle fibers (Rochlin et al. 2010). In the insect nervous system, for example, initially uniform neuroectodermal cells first invaginate, divide following a very defined pattern and eventually undergo morphological differentiation to give rise to highly polarized neurons with long axon projections and shorter dendrites (Skeath and Thor 2003; Reichert 2011). Other cell types, such as germ cells first migrate long distances before coming to rest in the developing gonads (Richardson and Lehmann 2010). Although these cell-type specific processes need to be tightly controlled and coordinated with those of other cell types of the same and neighboring organs, the molecular mechanisms involved are still poorly understood. The development of the adult *Drosophila melanogaster* head and the visual system has been proven to be an excellent model to study the coordination of different developmental processes (Atkins and Mardon 2009; María Domínguez and Casares 2005; Fernando Casares and Almudi 2016; Wolff and Ready 1991; J E Treisman and Heberlein 1998; Jessica E. Treisman 2013).

The adult *D. melanogaster* head is composed of the compound eyes (the main visual system), the three dorsal ocelli, the antennae, the ventral mouthparts and the head capsule that connects these organs and encloses the brain (Snodgrass 1935). Most of these structures develop during larval stages from eye-antennal imaginal discs, which originate from about 20 cells that are specified by *eyeless (ey)* expression at embryonic stages (Cohen 1993; Garcia-Bellido and Merriam 1969; Quiring et al. 1994). Throughout larval development, the eye-antennal discs grow extensively by cell proliferation resulting in discs composed of more than 15,000 cells at the beginning of pupation (Kenyon et al. 2003; Fernando Casares and Almudi 2016). During the first two larval stages, the initially uniform disc is subdivided into an anterior antennal and a posterior retinal compartment by the action of the two opposing gradients of the morphogens Wingless (Wg) and Decapentaplegic (Dpp), which subsequently activate genes responsible for antennal development (F Casares and Mann 1998; P. D. Dong, Chu, and Panganiban 2000) and the retinal determination genes (Cho et al. 2000; Chen et al. 1999; Cheyette et al. 1994; Kango-Singh, Singh, and Sun 2003; Mardon, Solomon, and Rubin 1994; Serikaku and O’Tousa 1994; Shen and Mardon 1997; Kenyon et al. 2003), respectively. Approximately at the same time when the retinal part of the disc and the antennal region separate during the early L2 stage, the ventral portion of the antennal part that gives rise to the maxillary palp is marked by expression of the Hox gene *Deformed (Dfd)*. This subdivision within the antennal region is established by delayed expression of *wg* in the Anlagen of the maxillary palps (Lebreton et al. 2008; V. K. Merrill et al. 1989; Anais Tiberghien et al. 2015).

Once the eye-antennal disc is subdivided into the different organ precursors, cells within each compartment start to differentiate at L2/early L3 stages. In the retinal region, a differentiation wave that is established in the posterior most part of the equator region moves anteriorly. This wave is accompanied by a morphologically visible indentation, the so-called morphogenetic furrow (MF) (Heberlein and Treisman 2000). Progression of the MF is mediated by Dpp-signaling within the furrow and Hh-signaling from the posterior disc margin (María Domínguez and Hafen 1997; Heberlein et al. 1995; J E Treisman and Heberlein 1998). Hh activated *atonal (ato)* expression in the region of the MF becomes restricted to regularly spaced single cells posterior to the furrow (M Domínguez 1999; María Domínguez and Hafen 1997). Those cells are destined to become R8 photoreceptors, which subsequently recruit R1-R7 photoreceptors and associated cell types, such as cone and pigment cells from the surrounding cells (N E Baker and Yu 2001; Jarman et al. 1994; Jarman et al. 1995).

The axons of successively forming photoreceptor cells need to be connected to the optic lobes to allow a functional wiring of the visual system with the brain. All axons are collected at the basal side of the eye-antennal disc and guided through the optic stalk throughout the L3 stage. This process is supported by retinal glia cells, which originate mainly by proliferation from 6-20 glia cells located in the optic stalk prior to photoreceptor differentiation (Silies et al. 2007; R Rangarajan, Gong, and Gaul 1999; Choi and Benzer 1994). These retinal glia cell types include migratory surface glia (including perineurial and subperineurial glia cells) and wrapping glia. Triggered by the presence of developing photoreceptor cells, the retinal glia cells enter the eye antennal disc through the optic stalk and migrate towards the anterior part of the disc, always remaining posterior to the advancing morphogenetic furrow (R Rangarajan, Gong, and Gaul 1999; Choi and Benzer 1994; Silies et al. 2007). When photoreceptors differentiate, the contact of their growing axons with perineurial glia cells triggers the reprogramming of these glia cells into differentiated wrapping glia, which extend their cell membranes to ensheath bundles of axons that project to the brain lobes through the optic stalk (Franzdóttir et al. 2009; Silies et al. 2007; Hummel et al. 2002). The basally migrating perineurial glia cells and the wrapping glia ensheathed projecting axons are separated by two large polyploid carpet cells, each of them covering half of the retinal field (Silies et al. 2007). The two carpet cells form septate junctions and express the G protein-coupled receptor (GPCR) encoded by the *moody* locus, both characteristics of the subperineurial surface glia type (Bainton et al. 2005; Silies et al. 2007). While subperineurial glia cells located in the brain remain there to form the blood-brain barrier, the carpet cells are thought to originate in the optic stalk (Choi and Benzer 1994), and during L2 and early L3 stages they migrate into the eye-antennal disc. Later during pupal stages, they migrate back through the optic stalk to remain beneath the lamina neuropil in the brain. However, so far it is not known, whether carpet cells or other retinal glia cell types eventually contribute to the formation of the blood-eye barrier, the retinal portion of the blood-brain barrier (T. N. Edwards et al. 2012; T. N. Edwards and Meinertzhagen 2010). The carpet cells thus share features of subperineurial glia, but their extensive migratory behavior and their function in the eye-antennal disc suggest that these cells may exhibit distinct cellular features. However, so far, no carpet cell specific regulator has been identified that may be involved in specifying carpet cell fate.

Although eye-antennal disc growth and patterning, and especially retinal determination and differentiation, are among the most extensively studied processes in *D. melanogaster*, a systematic understanding of involved genes and their potential genetic and direct interactions is limited to the late L3 stage in the context of retinal differentiation (Aerts et al. 2010; Naval-Sańchez et al. 2013; Potier et al. 2014). Similarly, recent attempts to incorporate existing functional and genetic data into a gene regulatory network context covers mainly retinal determination and differentiation processes (Koestler et al. 2015). So far, a comprehensive profiling of gene expression dynamics throughout eye-antennal disc development is missing. The same holds true for the molecular control of retinal glia cell development. While the transcriptome of adult surface glia in the brain has been analyzed (DeSalvo et al. 2014), retinal glia cells have not been comprehensively studied yet.

Here we present a dynamic genome wide expression analysis of *D. melanogaster* eye-antennal disc development covering late L2 to late L3 stages. We show that the transition from patterning to differentiation is accompanied by extensive remodeling of the transcriptional landscape. Furthermore, we identified central transcription factors that are likely to regulate a high number of co-expressed genes and thus key developmental processes in the different organ anlagen defined in the eye-antennal disc. One of these central factors is the C2H2 zinc-finger transcription factor Hunchback (Hb) (Tautz et al. 1987) that has been extensively studied in *D. melanogaster* during early axis determination and segmentation (Lehmann and Nüsslein-Volhard 1987; Nüsslein-Volhard and Wieschaus 1980). It is also well-known for its role in the regulation of temporal neuroblast identity during embryogenesis, as it determines first-born identity in the neural lineage (Grosskortenhaus et al. 2005; Isshiki et al. 2001). Here we show for the first time that *hb* is expressed in carpet cells and loss of function experiments suggest that its activity is necessary for carpet cell formation and consequently for proper axon guidance and blood-brain barrier integrity. Eventually, we reveal putative Hb target genes and confirm that bioinformatically predicted targets are indeed expressed in developing carpet cells.

## Results

### Differential gene expression and co-expressed genes during *D. melanogaster* head development

Although compound eye development and retinal differentiation are among the most intensively studied processes in *D. melanogaster*, a comprehensive understanding of the underlying gene expression dynamics is still missing to date. To identify the genes expressed during *D. melanogaster* eye-antennal disc development and their expression dynamics, we performed RNA-seq on this tissue at three larval stages covering the process of retinal differentiation that is marked by the progression of the morphogenetic furrow. The late L2 stage (72h after egg laying, AEL) represents the initiation of differentiation, at mid L3 stage (96h AEL) the morphogenetic furrow is in the middle of the retinal field and the late L3 stage (120h AEL) represents the end of morphogenetic furrow progression. Multidimensional scaling clustering clearly indicated that the largest difference in gene expression (dimension 1) was between L2 eye-antennal discs (72h AEL) and L3 eye-antennal discs (96h and 120h AEL) (Figure S1).

After filtering out not expressed and very lowly expressed genes, we observed that 9,194 genes were expressed at least in one of the three sequenced stages. As anticipated by the multidimensional scaling plot (Figure S1), the number of genes that changed their expression between 72h AEL and 96h AEL was much larger than between 96h AEL and 120h AEL, (Table S1). In only 24 hours, during the transition from L2 to L3, 60% of the expressed genes changed their expression significantly. In the transition from mid L3 to late L3, in contrast, only 22% of the genes underwent a change in their expression.

In order to better characterize the different expression dynamics of the expressed genes, we performed a co-expression clustering analysis based on Poisson Mixture models (Rau et al. 2015). Manual comparison of the different outputs showed that the 13 clusters predicted by one of the models (Djump, (Baudry et al. 2012)) were non-redundant and sufficiently described all the expression profiles present in the data. A total of 8,836 genes could be confidently placed in one of these clusters (maximum a posteriori probability (MAP) > 99%). We ordered the predicted 13 clusters according to their expression profile (Figure 1): four clusters contained clearly early expressed genes, two of them contained genes expressed only at 72h AEL (cluster 1 and 2) and two contained genes predominantly expressed early, but also with low expression at 96h and/or 120h AEL (clusters 3 and 4); one cluster showed down-regulation at 96h AEL, but a peak of expression again at 120h AEL (cluster 5); the genes in the largest clusters showed almost constant expression throughout the three stages (clusters 6 and 7); one cluster showed constant expression at 72h AEL and 96h AEL and down-regulation at 120h AEL (cluster 8); one cluster showed a peak of expression at 96h AEL (cluster 9) and four clusters contained genes with predominantly late expression, one with high and constant expression at 96h AEL and 120h AEL (cluster 10), two with up-regulation in both transitions (cluster 11 and cluster 12) and one with genes expressed only at 120h AEL (cluster 13).

**Figure 1.**
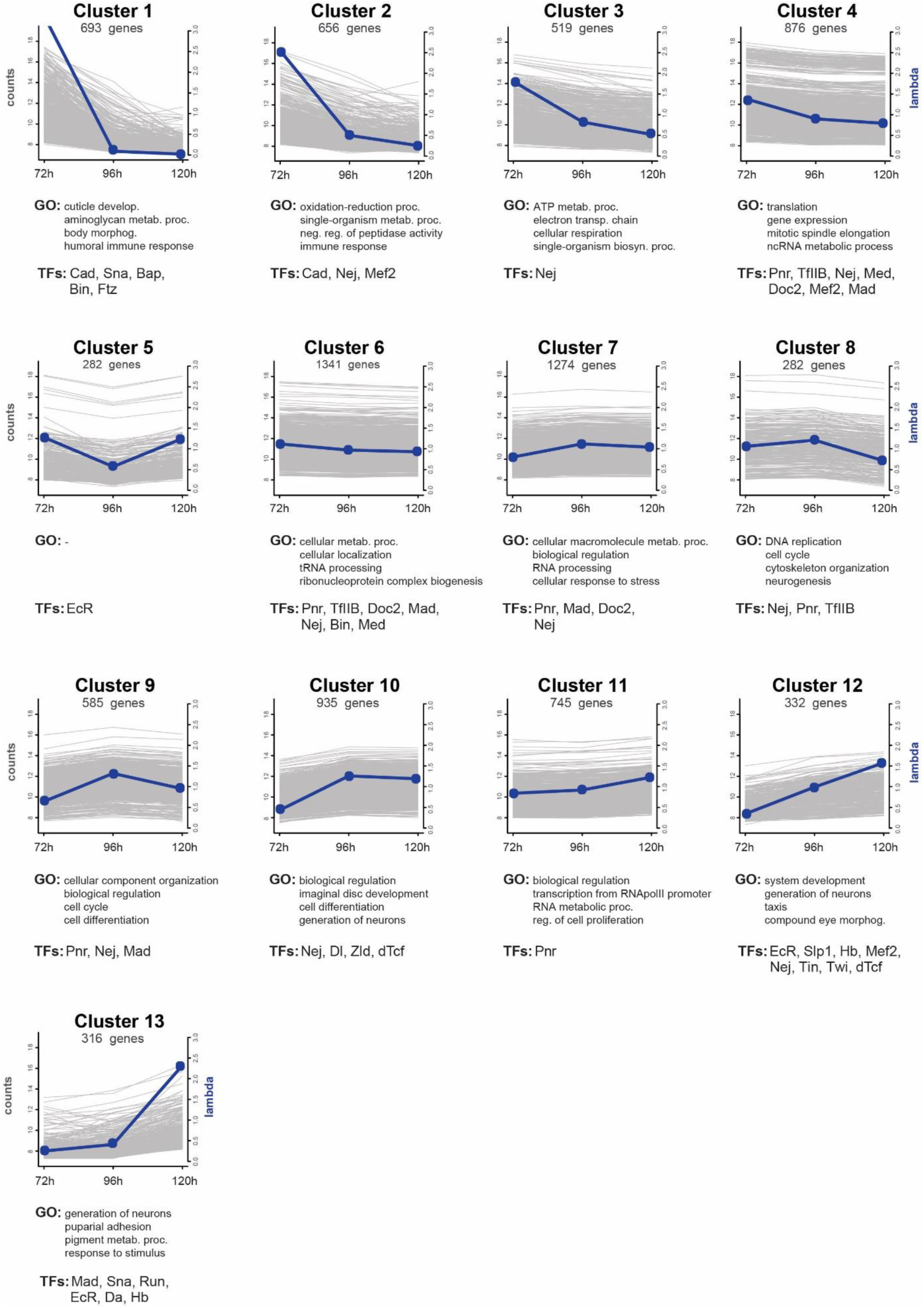
Co-expression clusters. All 13 profiles of co-expressed genes predicted by HTSCluster (see Materials and Methods). The number of genes assigned to a particular cluster are indicated below the cluster name. Blue dots represent relative expression levels (lambda value) of the genes of that cluster (y-axis on the right) at each stage. Background grey lines represent the normalized mean count of all genes belonging to a cluster (y-axis on the left). Below each cluster plot, the first four non-redundant GO terms enriched in the genes of that cluster are listed (see Table S2) and also the significantly enriched transcription factors (NES > 3, see Table S3).

A GO enrichment analysis for the genes in the individual clusters showed that the genome-wide co-expression profiling and subsequent ordering of the clusters recapitulated the consecutive biological processes that take place during eye-antennal disc development with a great resolution (Figure 1, Table S2). For instance, we found genes related to energy production mainly in clusters 2 and 3, while genes more specific for terms related to mitosis and cell cycle were found in clusters 4, 8 and 9, where genes have higher relative expression at 96h AEL than 72h AEL. Similarly, cluster 10 contained the more general term “imaginal disc development”, while cluster 12 showed enrichment for “compound eye morphogenesis”, and cluster 13 was the only with enriched terms related to pupation processes and pigmentation. Although we sequenced the entire eye-antennal discs, we found many GO terms related to eye development with high enrichment scores, while very few GO terms specific for antenna and maxillary palps were observed (e.g. in cluster 12 “eye development” appears with *p*=4.38e-24, “antennal development” with *p* =4.37e-08 and no GO terms related to maxillary palps were found) (Table S2). However, many GO terms related to leg formation and proximodistal pattern formation were highly enriched in the genes in cluster 9 (“proximaldistal pattern formation” with p=4.48e-05), cluster 10 (“leg disc development” with *p*=7.85e-20) and cluster 12 (“leg disc development” with *p*=4.32e-12) (Table S2). The assignment of all expressed genes to their corresponding cluster is available along with the GEO submission number GSE94915.

In summary, we showed that clusters with early expressed genes mainly represent metabolic and energy related processes, while clusters with late expressed genes represent more organ specific differentiation and morphogenetic processes.

### Transcription factors regulating *D. melanogaster* head development

The co-expression of genes observed in the 13 clusters may be a result of co-regulation by the same transcription factors or combinations thereof. In order to test this hypothesis and to reveal potential central upstream regulators, we used the i-cis Target method (Herrmann et al. 2012) to search for enrichment of transcription factor binding sites in the regulatory regions of the genes within each of the 13 clusters (Figure 1, Table S3). As basis for this enrichment analysis various experimental ChIP-chip and ChIP-seq datasets were used, namely those published by the modENCODE Consortium (Celniker et al. 2009), the Berkeley *Drosophila* Transcription Network Project (X. Li et al. 2008) and by the Furlong Lab (Zinzen et al. 2009; Junion et al. 2012).

One of the most noticeable results of the statistical ranking analysis was that genes in 9 of the 13 clusters showed significant enrichment for Nejire binding sites (Figure 1). Nejire (also known as CREB-binding protein (CBP)) is a co-factor already known to be involved in many processes of eye development and patterning (Justin P Kumar et al. 2004). Similarly, Pannier that has been shown to play at least two important roles during eye-antennal disc development (Singh et al. 2005; Singh and Choi 2003; Oros et al. 2010) was found enriched to regulate the genes of many clusters (clusters 2, 4, 6, 7, 8, 9 and 11).

Besides these highly abundant transcription factors, the clusters with genes predominantly expressed at later stages were also enriched for transcription factors already known to play a role in eye-antennal disc development. For instance, a significant number of Sloppy-paired 1 (Slp1) target genes are up-regulated at L3 stage (cluster 12) and this transcription factor is known to play a critical role in establishing dorsal-ventral patterning of the eye field in the eye-antennal disc (Sato and Tomlinson 2007). A function of Daughterless (identified in cluster 13) is also described: it is expressed in the morphogenetic furrow, it interacts with Atonal and is necessary for proper photoreceptor differentiation (Brown et al. 1996). Finally, Snail (enriched in cluster 1 and 13) and Twist (enriched in cluster 12) were previously identified as possible repressors of the retinal determination gene *dachshund (dac)* (Anderson, Salzer, and Kumar 2006) and our results could indicate that they regulate also other genes during eye-antennal disc development.

Cluster 5 contained genes that show a peak in expression at 72h AEL and 120h AEL stages, which precede major stage transitions from L2 to L3 and from L3 to pupa stage, respectively. These transitions are characterized by ecdysone hormone pulses before larval molting and pupation (T. Li and Bender 2000). Intriguingly, the only potential transcription factor binding site that was significantly enriched was that of the Ecdysone Receptor (EcR), that has been shown to be expressed in the eye-antennal disc in the region of the progressing morphogenetic furrow (Brennan et al. 1998).

The identification of many well-known transcription factors suggests that the applied clustering approach indeed allows identifying key regulators of various processes taking place throughout eye-antennal disc development. Interestingly, we identified a few generally well-known upstream factors for which a potential role during eye-antennal disc development has not yet been described. For instance, in clusters of very early expressed genes, we found an enrichment of motifs for the transcription factor Caudal (Cad) (cluster 1 and 2) and the Hox protein Fushi tarazu (Ftz) (cluster 1). The MADS-box transcription factor Myocyte enhancer factor 2 (DMef2) was predicted to regulate genes found in clusters 2, 4 and 12 (Figure 1). Using two independent *Dmef2*-Gal4 lines to drive GFP expression, we confirmed expression of *Dmef2* in lose cells attached to the developing eye-antennal discs (Figure S2). Eventually, we found an enrichment of potential target genes of the C2H2 zinc-finger transcription factor Hunchback (Hb) in clusters 12 and 13, which are active mainly during mid and late L3 stages. Since GO terms enriched in these two clusters suggested an involvement in retinal development or neurogenesis (Figure 1), we examined a potential function of Hb in the eye-antennal disc in more detail.

### *hb* is expressed in retinal subperineurial glia cells

Using *in-situ* hybridization we found *hb* expression in two cell nuclei at the base of the optic stalk in the posterior region of late L3 eye-antennal discs (Figure 2A). With a Hb antibody we also detected the Hb protein in these two basally located nuclei (Figure 2B). DNA staining with DAPI showed that the Hb-positive nuclei are bigger than those of surrounding cells, suggesting that they are polyploid. Additionally, we tested two putative Gal4 driver lines obtained from the Vienna Tile library (Pfeiffer et al. 2008) (VT038544; Figure 2C and VT038545; Figure S3). Both lines drove reporter gene expression in the two polyploid nuclei as described above. Note that both lines also drove the typical *hb* expression in the developing embryonic nervous system, but not the early anterior expression (Jiménez and Campos-Ortega 1990; Kambadur et al. 1998) (not shown). The regulatory region covered by the two Gal4 driver lines is located at the non-coding 3’ end of the *hb* locus accessible to DNA-binding proteins at embryonic stages 9 and 10 (X. Li et al. 2008) (Figure S4), a time when early-born neuroblasts express *hb* (Grosskortenhaus et al. 2005). The lack of the early anterior expression may be explained by the fact that the DNA region covered by the driver lines does not seem to be bound by Bicoid during early embryonic stages (X. Li et al. 2008) (Figure S4). Based on these findings, we are confident that the regions covered by the two Gal4 driver lines (VT038544 and VT038545) recapitulate native *hb* expression.

**Figure 2.**
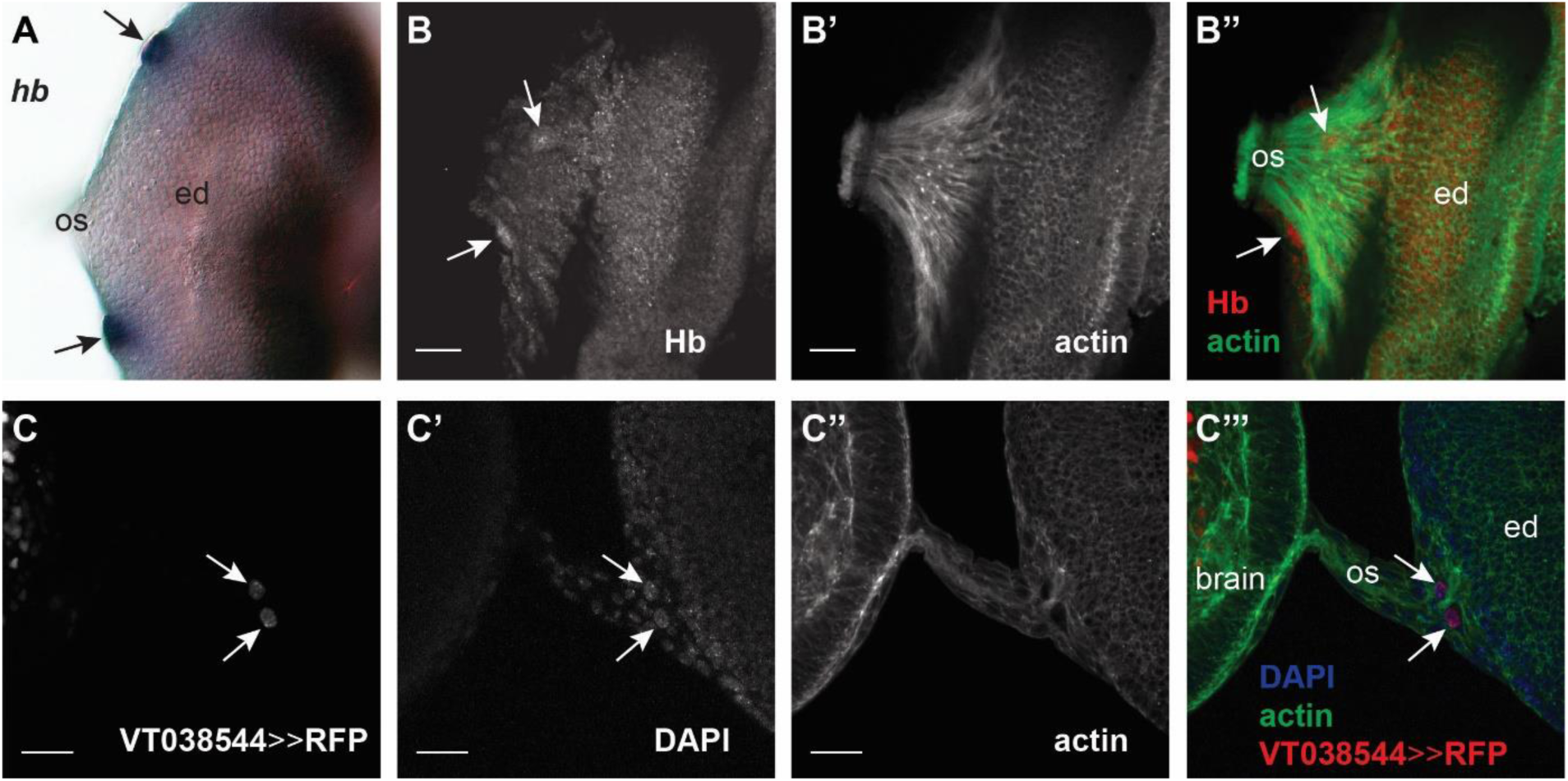
*hb* expression in the eye-antennal disc. Different detection methods showing *hb* expression in L3 eye-antennal discs. In all pictures, anterior is to the right. Eye disc (ed), optic stalk (os). Scale bar = 20 μm. **A.** *in-situ* hybridization of *hb* mRNA. *hb* is expressed in two large domains (black arrows) at the posterior end of the eye field **B.** Antibody staining of Hb protein (rabbit α-Hb), showing a signal in two large domains (white arrows) at the posterior end of the eye field. Co-staining with Phalloidin (**B’**, **B”**) shows that the cells expressiong *hb* are located between the photoreceptor axons on their way to the optic stalk. **C.** Expression of histone-bound RFP (UAS-H2B::RFP) driven by a Gal4 line containing an enhancer region near the *hb* locus (VT038544-Gal4 driver line obtained from the Vienna Tile collection, see Figure S4 for details). Expression is localized in two large cells (**C’**) (white arrows) in the same location as A and B, at the posterior end of the eye field, near the optic stalk (**C’’’**).

The basal location of the *hb*-positive cells suggests that they may be retinal glia cells. Coexpression of *hb* with the pan-glial marker Repo (Figure 3A) further supported this suggestion. Previous data has shown that two polyploid retinal subperineurial glia cells (also referred to as carpet cells) cover the posterior region of the eye-antennal disc (Choi and Benzer 1994; Silies et al. 2007). In order to test, whether *hb* may be expressed in carpet cells, we first investigated the expression of the subperineurial glia marker Moody (Schwabe et al. 2005) and we found a clear co-localization with Hb (Figure 3B).

**Figure 3.**
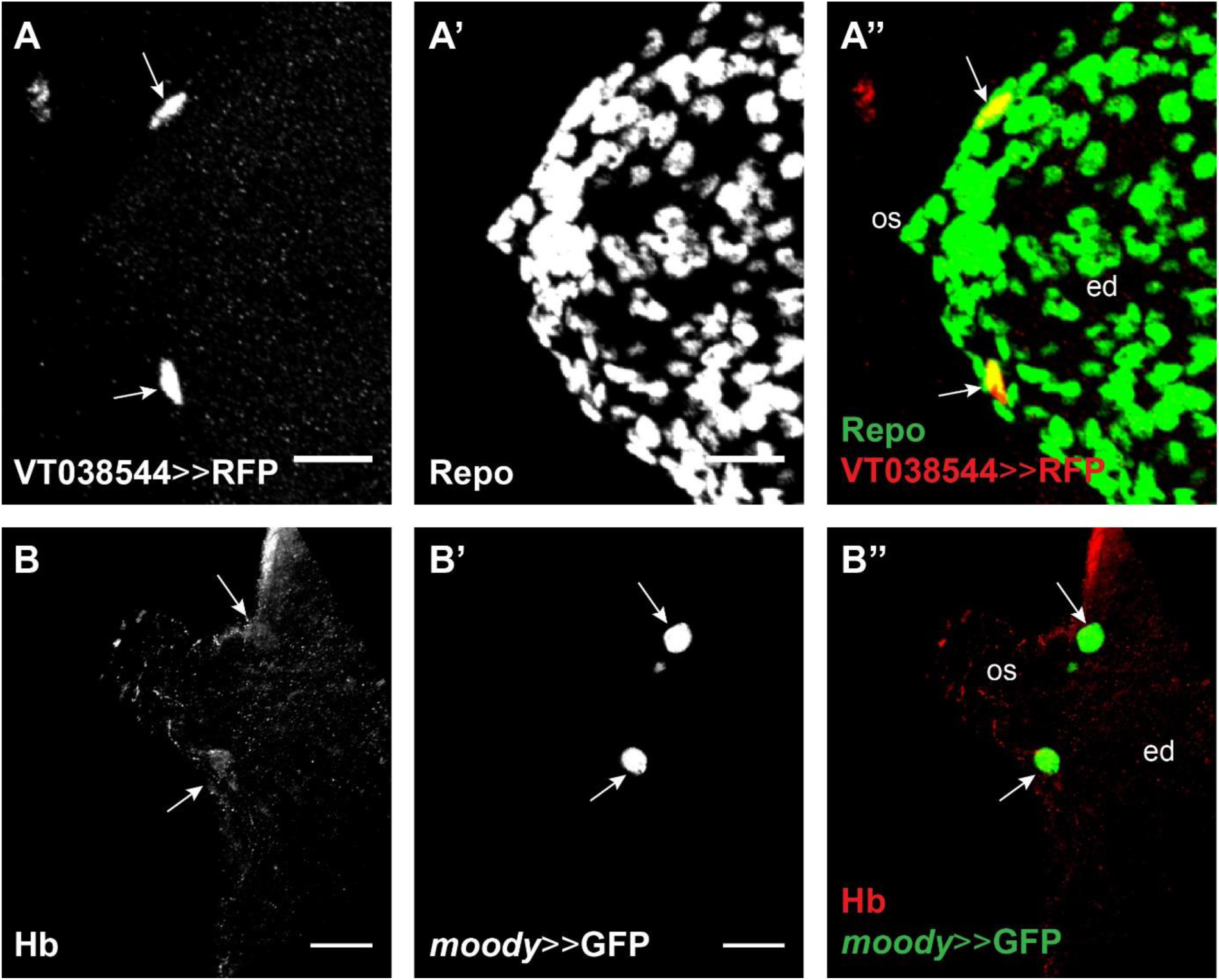
*hb* is expressed in subperineurial glia cells. **A.** *hb* expression (VT038544-Gal4 driving UAS-GFP, green) co-localizes (white arrows) with the pan-glial marker Repo (detected with rabbit α-Repo antibody, red). **B.** Hb (detected with rabbit α-Hb antibody, red) co-localizes (white arrows) with the expression of the subperineurial glia cell marker *moody* (*moody*-Gal4 driving UAS-GFP, green). In all pictures, anterior is to the right. Eye disc (ed), optic stalk (os). Scale bar = 20 μm.

Carpet cells migrate through the optic stalk into the eye-antennal disc during larval development (Choi and Benzer 1994; Silies et al. 2007). Therefore, we followed the expression of the *hb* driver lines throughout late L2 and L3 larval stages (Figure 4). Already at the L2 stage, we could easily recognize the *hb*-positive cell nuclei by their large size (Figure 4A, A’). We could corroborate that these cells indeed migrated through the optic stalk during late L2 and early L3 stages (Figure 4A, B), and then entered the disc and remained basally in the posterior region of the disc, flanking the optic stalk (Figure 4C, C’’). As previously observed for carpet cells (Silies et al. 2007), we never found *hb*-positive cell nuclei in the midline of the retinal field.

**Figure 4.**
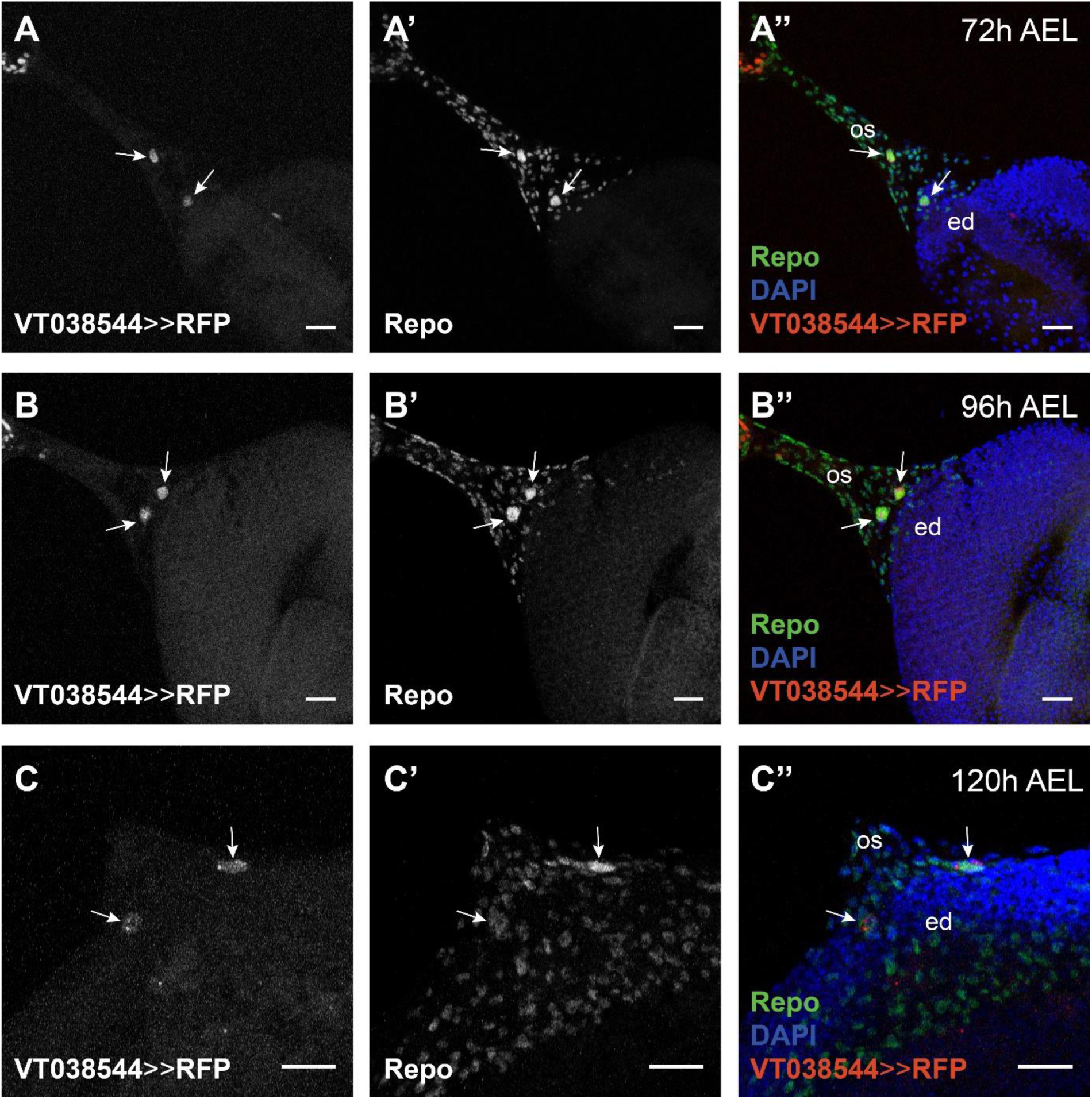
*hb* is expressed in cells that migrate from the optic stalk into the eye-antennal discs. *hb*-expressing cells are visualized with VT038544-Gal4 driving histone-bound RFP (UAS-H2B::RFP, red) (**A**, **B** and **C**) and all glia cells are stained with rabbit α-Repo antibody (**A’**, **B’** and **C’**). Eye disc (ed), optic stalk (os). Scale bar = 20 μm. **A.** During L2 stage, the glia cells expressing *hb* are located in the optic stalk (white arrows). **B.** At mid L3 stage, these cells are located at the edge between the optic stalk and the retinal region of the eye-antennal disc (white arrows). **C.** At late L3 stage, these cells are located in the posterior region of the eye field, on at each side of the optic stalk (white arrows).

Taken together, these data show that *hb* is expressed in two polyploid retinal subperineurial glia cells (carpet cells) that enter the basal surface of the eye-antennal disc through the optic stalk during larval development.

### Hb function is necessary for the presence of polyploid carpet cells in the eye-antennal disc

The expression of *hb* in carpet cells suggested an involvement in their development. To test this hypothesis, we examined loss of Hb function phenotypes based on RNA interference (RNAi) driven specifically in subperineurial glia cells (*moody*-Gal4 driving UAS-*hb*^dsRNA^). Of four tested UAS-*hb*^dsRNA^ lines we used the most efficient line (see Materials and Methods) for the RNAi knockdown experiments. Additionally, we investigated eye-antennal discs of a temperature sensitive mutant (Hb^TS^) (Bender, Turner, and Kaufman 1987). Since Hb is necessary during embryogenesis (Nüsslein-Volhard and Wieschaus 1980; Lehmann and Nüsslein-Volhard 1987), the analyzed flies were kept at 18°C during egg collection and throughout embryonic development, and they were only transferred to the restrictive temperature of 28°C at the L1 stage. Carpet cell nuclei were identified by α-Repo staining because of their large size and their specific position (Figure 5A; see also Figure 4B’ and 4C’).

**Figure 5.**
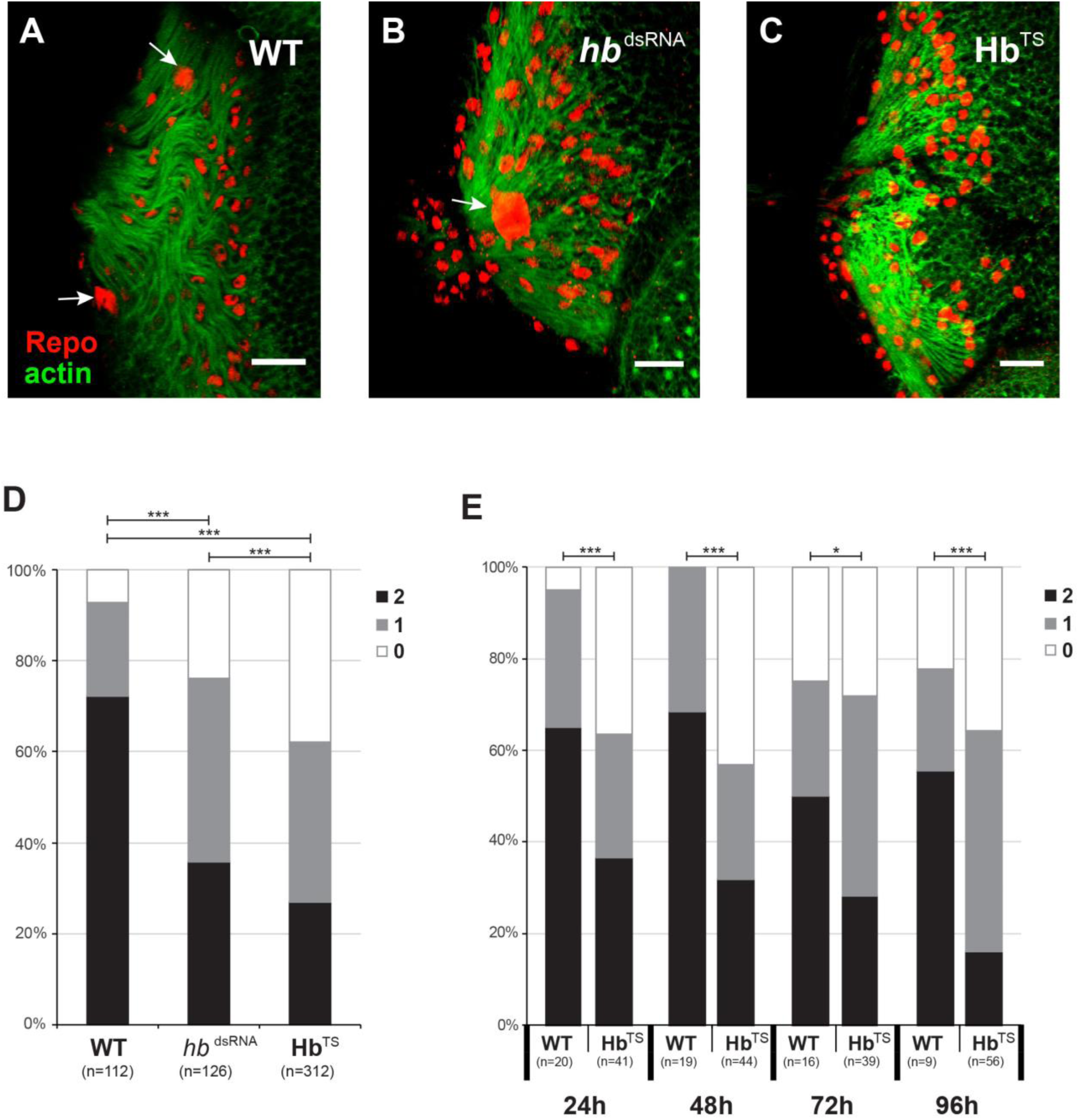
The number of polyploid glia cell nuclei is reduced after loss of Hb function. **A-C.** Staining with rabbit α-Repo antibody (red) and Phalloidin (green) of late L3 eye-antennal discs in wild type (**A**), *repo* driven *hb* RNAi (**B**) and Hb temperature sensitive mutant (**C**). This figure represent the phenotypes that have been analyzed in **D**, where the number of polyploid nuclei (white arrows) have been quantified. In all pictures, anterior is to the right. Scale bar = 20 μm. **D.** Quantification of the number of polyploid nuclei observed in wild type (WT), *repo*-Gal4 and *moody*-Gal4 driven UAS-*hb* RNAi (*hb*^dsRNA^) and Hb temperature sensitive mutant (Hb^TS^). **E.** Quantification of the number of polyploid nuclei observed in late L3 eye-antennal discs of flies that have been raised at 18°C until the indicated time points (24h AEL, 48h AEL, 72h AEL and 96h AEL), when they have been transferred to the restrictive temperature of 28°C. In **D** and **E**, the black bar indicates percentage of discs with two polyploid glia cell nuclei, grey indicates discs with one polyploid glia cell nucleus and white indicates discs without polyploid glia cell nuclei. Pearson’s Chi-squared test was performed to determine if the distribution of the different number of cells (0, 1 or 2) was equal between wild type and RNAi. *: p-val < 0.05, ***: p-val < 0.0005.

The most common phenotype observed in late L3 eye-antennal discs of RNAi and mutant flies was the absence of one or both carpet cell nuclei (Figure 5A-C). In wild type animals, we could unambiguously identify two carpet cell nuclei in 72% of the eye-antennal discs. In 21% of the analyzed discs, we found only one carpet cell nucleus (Figure 5D). In contrast, in 35% to 40% of the studied Hb loss of function discs only one carpet cell nucleus was observed (Figure 5B and D). In some cases, this single polyploid Repo-positive nucleus was located in the midline of the retinal field (Figure 5B). No carpet cell nuclei could be observed in 24% and 38% of the eye antennal discs originating from *moody*>>*hb*^dsRNA^ and Hb^TS^ flies, respectively (Figure 5C and D). Note that we obtained comparable results when we expressed the *hb* dsRNA in all glia cells (*repo*>>*hb*^dsRNA^; not shown) or only in subperineurial glia cells (*moody*>>*hb*^dsRNA^).

To identify larval stages at which Hb function is crucial for carpet cell development, we transferred Hb^TS^ flies to the restrictive temperature of 28°C at 24h AEL (early L1 stage), at 48h AEL (late L1), at 72h AEL (late L2) or 96h AEL (mid L3 stage) and assessed the presence of polyploid Repo-positive carpet cell nuclei in late L3 eye-antennal discs, respectively. In all cases, we found a significant reduction of the number of carpet cell nuclei when compared to control discs (Figure 5E). Although no clear significant differences in the number of carpet cells was detected between the consecutive experiments (Figure S5), our results show that Hb function is necessary for the presence of polyploid carpet cell nuclei throughout larval development.

### Loss of Hb function affects retinal glia cell migration and axon guidance

The observed loss of polyploid carpet cell nuclei could be a result of either the loss of the entire carpet cells, incomplete migration into the eye-antennal disc or loss of the polyploidy. To distinguish between these options, we tested whether also the carpet cell membranes were affected upon loss of *hb* expression, in addition to the polyploid nuclei. To this aim, we expressed *hb_dsRNA_* specifically in subperineurial glia cells with a *moody*-Gal4 driver line together with a strong membrane marker (20xUAS-mCD8::GFP) to label the extensive carpet cell membranes (Figure 6).

**Figure 6.**
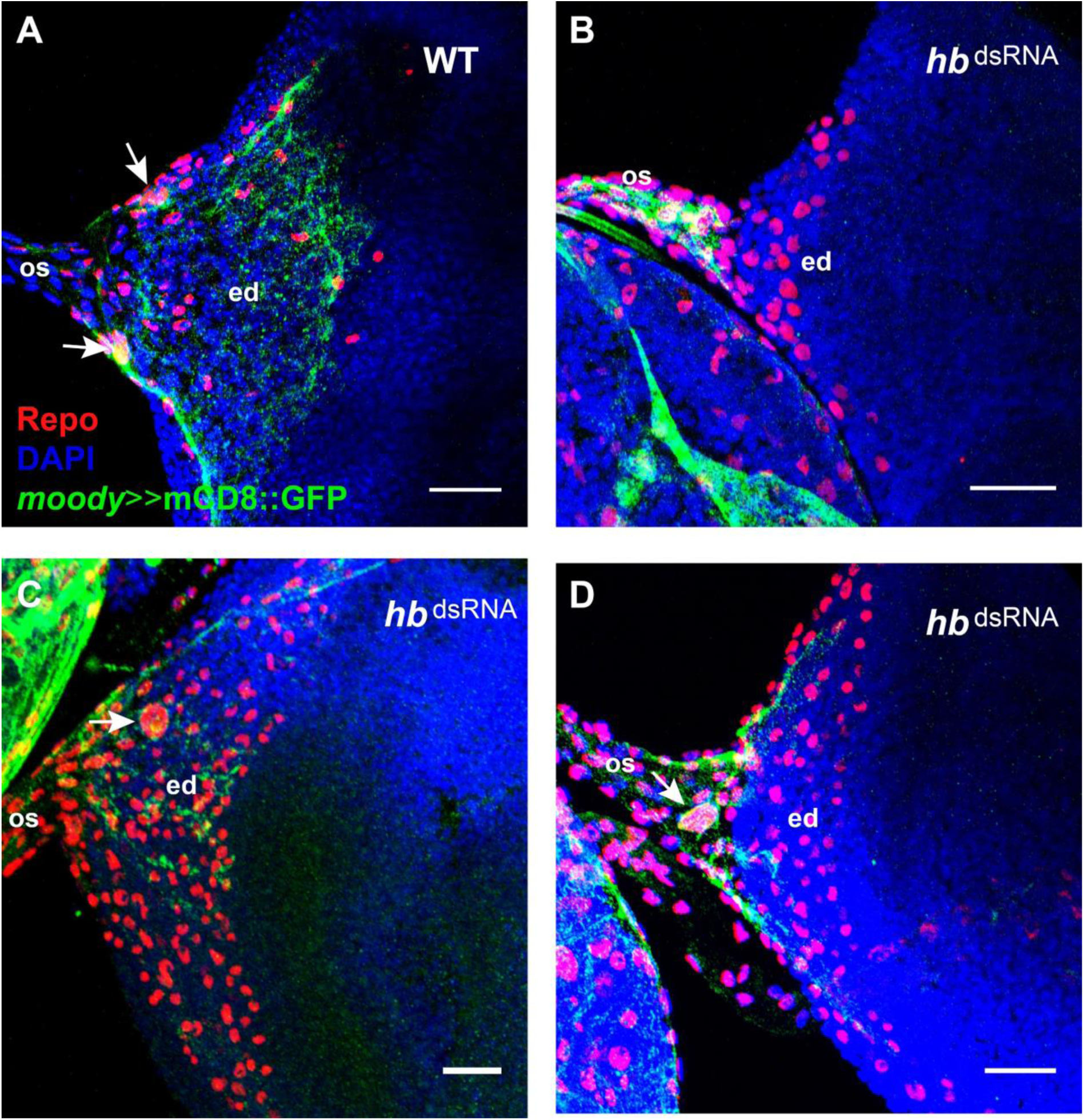
Carpet cell membranes after loss of Hb function. Membranes of carpet cells in late L3 eye-antennal discs are labelled with *moody*-Gal4 driven UAS-mCD8::GFP expression (green). All glia cells are stained with rabbit α-Repo antibody (red). Carpet cell nuclei (white arrows) are recognized by their large size. In all pictures, anterior is to the right. Eye disc (ed), optic stalk (os). Scale bar = 20 μm. **A.** In wild type, the membranes of the two carpet cells cover all the retinal field up to the edge of the most anteriorly located glia cells. **B-D.** Phenotypes observed after *moody* driven *hb* RNAi. In discs where carpet cell nuclei cannot be observed, GFP signal is detected only in the optic stalk (**B**). In discs where only one carpet cell nucleus can be observed on one side, the membrane signal is predominantly observed on that side (**C**). In discs where only one carpet cell can be observed in the disc midline, membrane extend to both sides (**D**), but do not extend so far anteriorly as in wild type (compare **D** to **A**).

In control discs (*moody*>>20xmCD8::GFP) the two carpet cell membranes spanned the entire posterior region of the eye-antennal disc from the optic stalk to the morphogenetic furrow (Figure 6A). In contrast, in some of the knock-down (*moody*>>20xmCD8::GFP; *moody*>>*hb*^dsRNA^) eye antennal discs with no clear polyploid carpet cell nuclei, we detected *moody*-positive membranes that remained in the optic stalk and did not span the entire retinal field of the eye-antennal disc (Figure 6B). In cases where one clear carpet cell nucleus was observed, the location of *moody*-positive cell membranes in eye-antennal discs depended on the location of the remaining nucleus. If the nucleus was located on one side of the eye-antennal disc, we observed *moody*-positive membranes more unilaterally (Figure 6C), while the membrane was present in the center of the disc if the polyploid nucleus was located centrally (Figure 6D).

It has been shown that the extensive cell bodies of carpet cells provide a scaffold for other retinal glia cells that migrate into the eye-antennal disc, pick up differentiating photoreceptor axons and guide them through the optic stalk into the optic lobe (Choi and Benzer 1994; R Rangarajan, Gong, and Gaul 1999). In accordance with this known function, we observed irregular and patchy patterns of Repo-positive cells in late L3 Hb loss of function eye-antennal discs, suggesting impaired glia cell migration into the eye-antennal disc (compare Figure S6B to S6A). Additionally, we used HRP staining to visualize axon projections in late L3 eye-antennal discs. While axonal tracts were regular in control eye-antennal discs, we found unorganized axon projections upon loss of *hb* expression (compare Figure S6B’ to S6A’).

### Loss of Hb function results in blood-brain barrier defects

Subperineurial glia cells cover the entire surface of the brain from larval stages onwards. They are an integral part of the protective blood-brain barrier by establishing intercellular septate junctions (Carlson et al. 2000). The blood-brain barrier prevents the substances that circulate in the hemolymph to enter the brain and helps maintaining the proper homeostatic conditions of the nervous system (J. S. Edwards, Swales, and Bate 1993). Since it has been shown that the carpet cells migrate through the optic stalk towards the brain during pupal stages (T. N. Edwards et al. 2012), we tested, whether the loss of *hb* expression in developing carpet cells had an effect on the integrity of the blood-brain barrier.

To this aim, we injected fluorescently labeled dextran into the abdomen of *moody*>>*hb*^dsRNA^ adult flies and scored the presence of this dye in the retina of the flies. Animals with a properly formed blood-brain barrier showed a fluorescent signal in their body, but not in the retina (Figure 7A). However, in animals that had an incomplete blood-brain barrier, the dextran penetrated into the retina and fluorescence was observed in the compound eyes (Figure 7A’). Since it is known that blood-brain barrier permeability can increase after exposure to stress conditions (H. S. Sharma and Dey 1986; Skultétyová, Tokarev, and Jezová 1998), we only scored animals that survived 24h after the injection of dextran. In most cases, the two eyes of an individual presented different fluorescent intensities, and even no fluorescence in one eye but strong signal in the other. Therefore, we scored each eye separately. *moody*>>*hb*^dsRNA^ flies had a significantly higher rate of fluorescent retinas (*p* = 8.08e-7, X^2^ test), indicating that their eyes were not properly isolated from the hemolymph circulating in the body cavity (Figure 7B).

**Figure 7.**
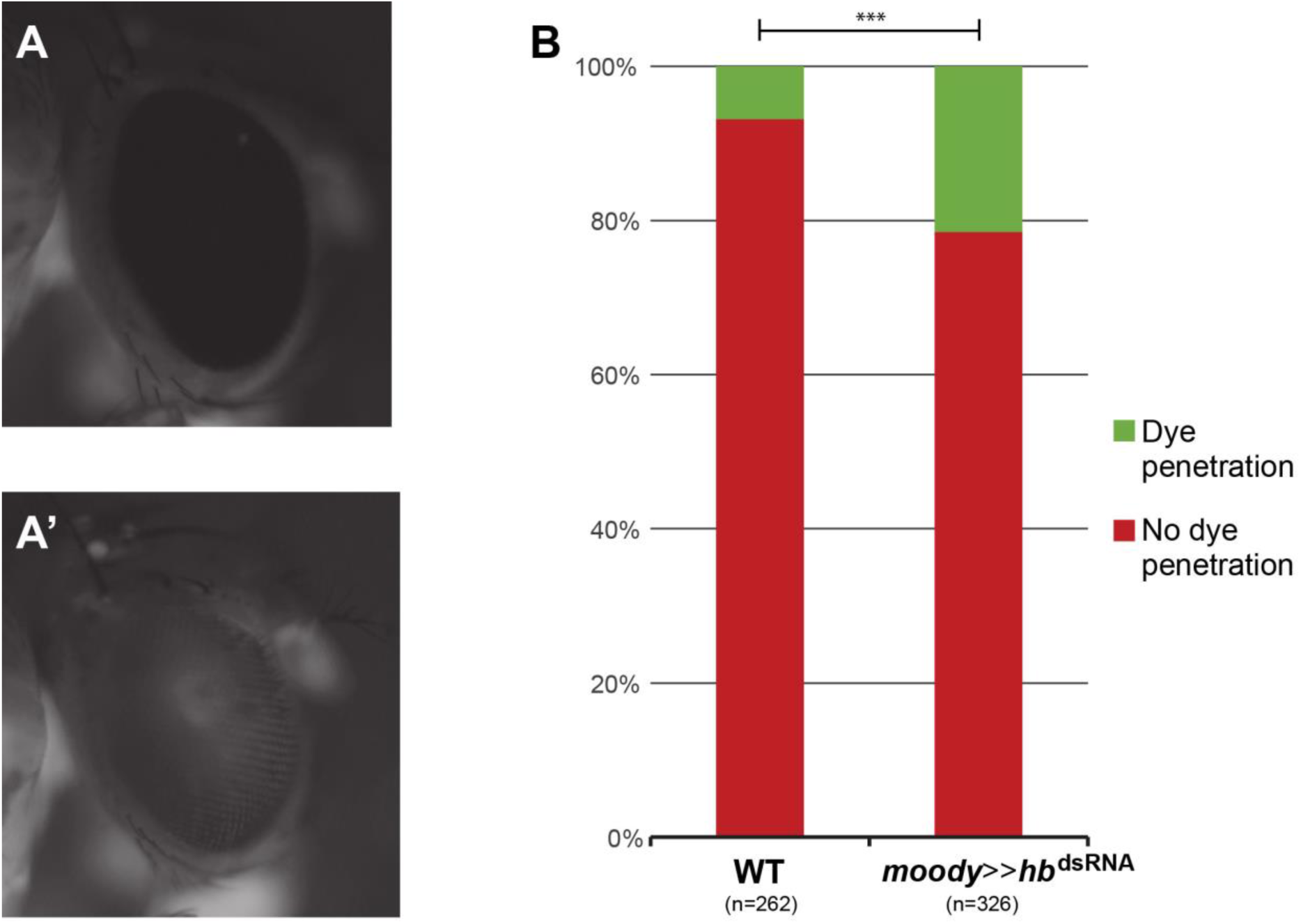
Blood-brain barrier function is impaired after loss of *hb* expression in carpet cells. **A.** After injection of fluorescently labelled dextran in the abdomen of adult flies, animals with correctly formed blood-brain barrier present fluorescence in the body (not shown) but not in the compound eye. **A’.** In flies with incomplete blood-brain barrier, fluorescent dye can be observed in the compound eye as well as in the body. **B.** Quantification of eyes with (green) or without (red) dye penetration. *hb* knock-down flies have a significant increase in the penetrance of dye in the eye, indicating a defective blood-eye barrier. Pearson’s Chi-squared test was performed to determine significance between the wild type results and the RNAi. ***: p-val < 0.0005.

In summary, our loss of function experiments further confirmed a central role of Hb in carpet cell development. Besides impaired retinal glia cell migration and axon guidance, we showed that upon loss of Hb function also the blood-brain barrier integrity is disrupted.

### Expression of putative Hb target genes in eye-antennal discs

Since we have identified Hb because of an increase in expression of its target genes during 96h and 120h AEL stages and *hb* itself is only expressed in carpet cells, we also investigated, whether some of the targets were expressed in these cells. Using available ChIP-chip data for Hb from the Berkeley *Drosophila* Transcription Network Project (BDTNP) (X. Li et al. 2008), we generated a high confidence list of 847 putative Hb target genes (see Materials and Methods for details), of which 585 were expressed in eye-antennal discs at least in one of the studied stages. More precisely, we found that 267 of these genes were differentially expressed in the transition from 72h to 96h AEL and only 52 were differentially expressed between 96h and 120h AEL (Figure 8, Table S4). In both cases, most of these genes were up-regulated, suggesting that Hb mainly activates target gene expression in the eye-antennal disc. Focusing only on those target genes that resulted in the identification of Hb in our clustering approach (see above), we found that 77 of the 585 expressed putative Hb targets were present in clusters 12 and 13. We searched the GO terms for biological functions of these 77 genes and found that 17 code for transcription factors and up to 25 code for proteins integral to the cell membrane. A number of GO terms were related to neuronal development and eye development and to note is the presence of genes known to be related to glia cell migration and endoreduplication (Table S5).

**Figure 8.**
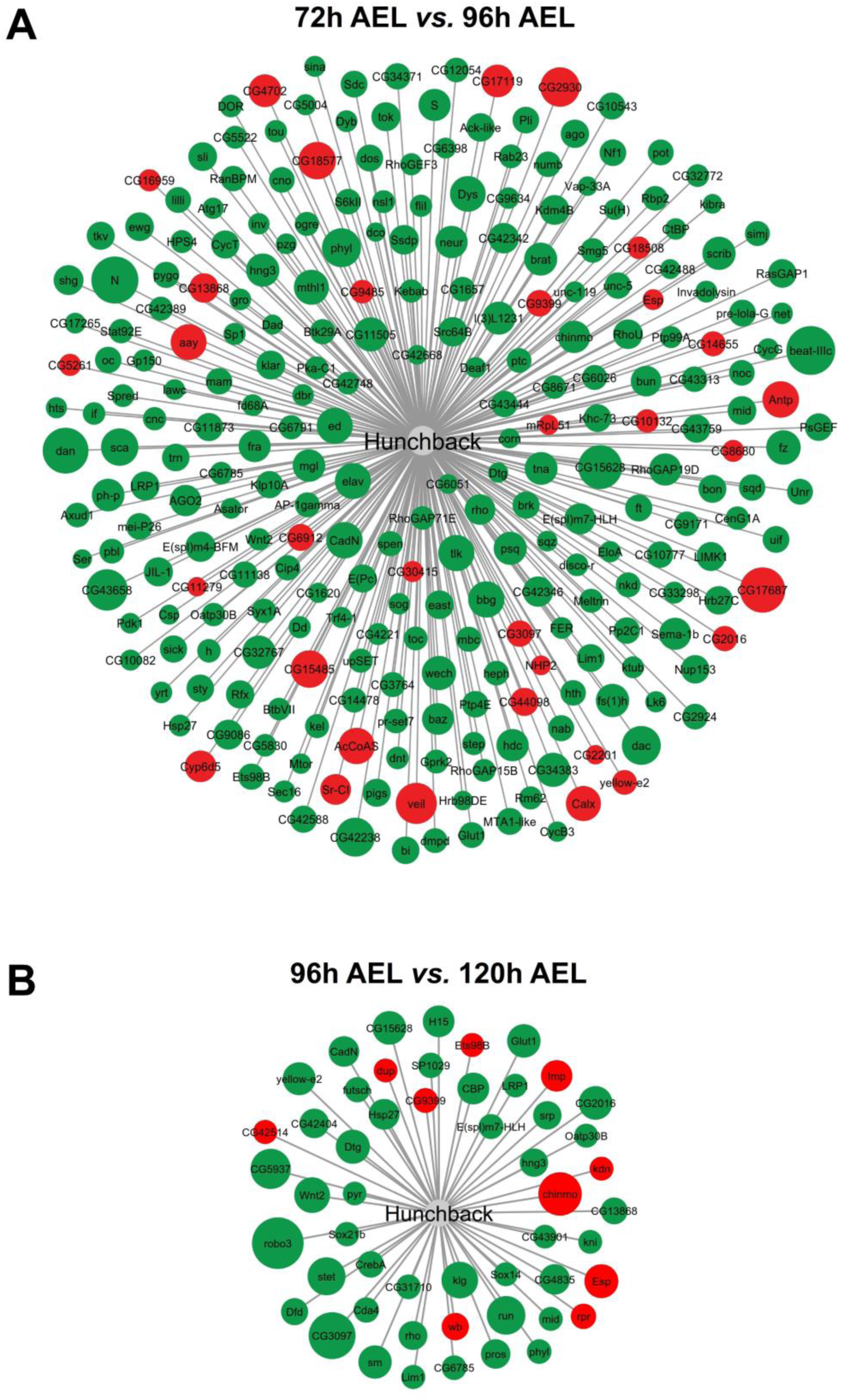
Differentially expressed putative Hb target genes. Green and red shaded circles are up- and down-regulated genes, respectively. Larger circle size indicates higher log2-fold change. **A.** 267 genes from the high confidence list of Hb targets are differentially expressed in the eye-antennal discs during the transition from late L2 (72h AEL) to mid L3 (96h AEL) stages. 33 genes are down-regulated and 234 are up-regulated (see Table S4). **B.** 52 genes from the high confidence list of Hb targets are differentially expressed in the eye-antennal discs during the transition from mid L3 (96h AEL) to late L3 (120h AEL) stages. 10 genes are down-regulated and 42 are up-regulated (see Table S4).

Based on their annotated GO terms, predicted or known cellular location and the availability of driver lines and antibodies, we selected 13 of these target genes and tested if they were expressed in carpet cells at 120h AEL. For 8 out of the 13 selected targets we found no clear expression related to carpet cells (*archipelago* (*ago*), *Delta* (*Dl*), *knirps* (*kni*), *rhomboid* (*rho*), *roundabout 3* (*robo3*), *Sox21b*, *Src oncogene at 64B* (*Src64B*) and *thickveins* (*tkv*), not shown). This could be because they were false positives, but they could also be expressed at earlier stages than analyzed here or the used driver constructs did not include the regulatory regions to drive expression in carpet cells. *brinker* (*brk*), *Cadherin-N* (*CadN*), *cut* (*ct*), *Fasciclin 2* (Fas2) and *sprouty* (*sty*) showed expression in carpet cells (Figure 9). *brinker* (*brk*) was ubiquitously expressed in the eye-antennal disc (not shown). Although we could only observe expression in one of the two cells in every eye-antennal disc we analyzed, *CadN* is clearly expressed in carpet cells (Figure 9A). Recent data demonstrated that CadN, a Ca^+^ dependent cell adhesion molecule, is necessary for the proper collective migration of glia cells (A. Kumar et al. 2015), a key feature of carpet cells. As it has previously been published, *cut* is expressed in subperineurial glia cells (Figure 9B) (Bauke et al. 2015). The Cut protein is present in carpet cells already at L2 stage and remains until late L3 stage (Figure 9B, earlier stages not shown). It has been shown that Cut is necessary for proper wrapping glia differentiation and to correctly form the large membrane processes that these cells form (Bauke et al. 2015). Interestingly, carpet cells have a similar morphology, with very large membrane surface and extensive processes that reach to the edge of the retinal field. In contrast, retinal perineurial glia do not have this morphology and do not express *cut*. Also, *Fas2* (Figure 9C) and *sty* (Figure 9D) were clearly expressed in carpet cells as well as in several other cells in the eye-antennal disc. Sty and Fas2 are negative regulators of the EGFR signaling pathway that is involved in retinal glia cell development and photoreceptor differentiation (Sieglitz et al. 2013; Jarvis 2006; Bogdan and Klämbt 2001; Kim and Bar-Sagi 2004; Kramer et al. 1999; Mao and Freeman 2009).

**Figure 9.**
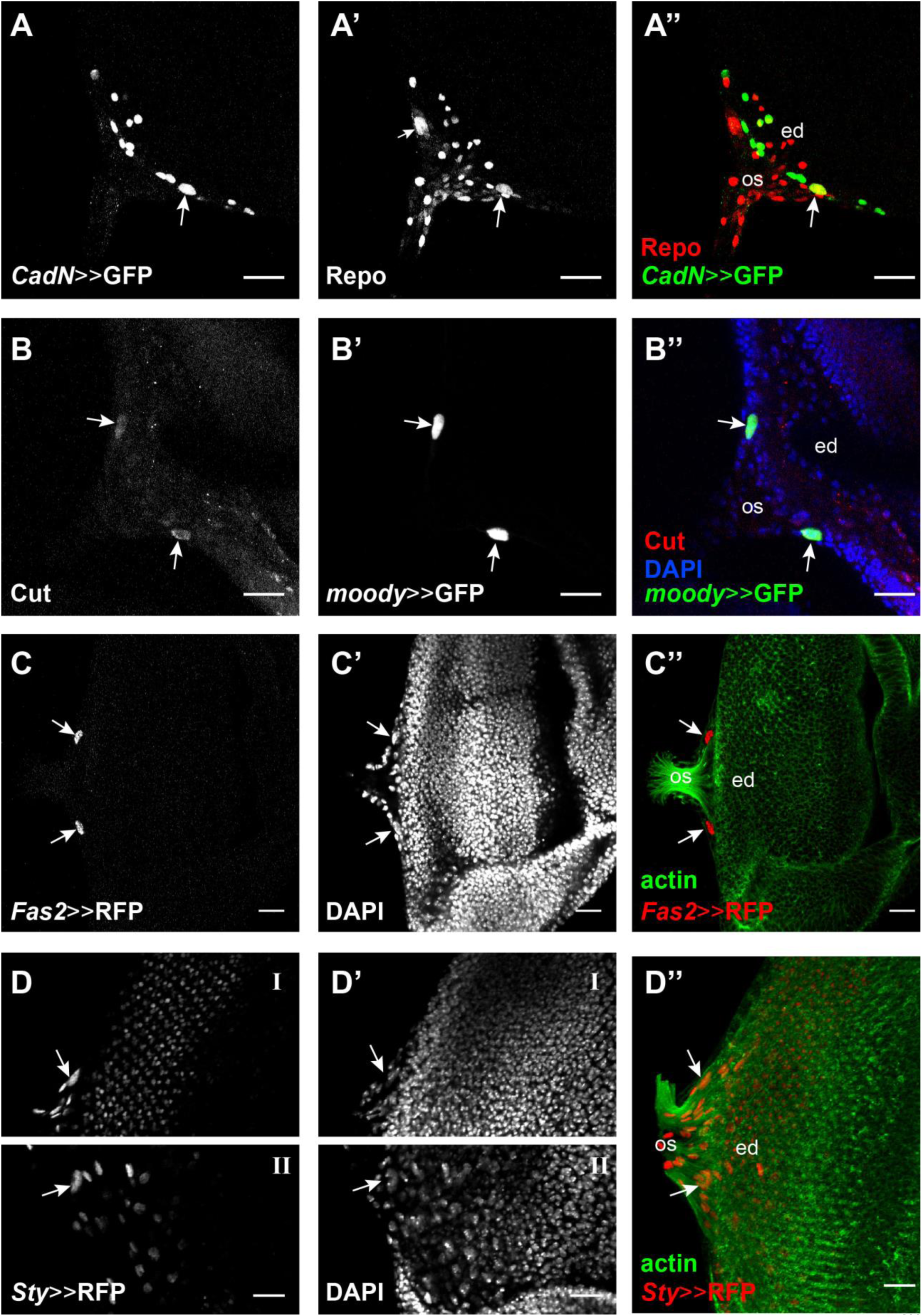
Expression of Hb target genes in the eye-antennal discs. Four of the tested target genes show expression in carpet cells. Eye disc (ed), optic stalk (os). Scale bar = 20 μm. **A.** *CadN*-Gal4 drives UAS-GFP expression (green in **A”**) in one of the two carpet cells (white arrow), as well as other cells in the disc, possibly glia cells. **A’** and **A”.** Carpet cells are recognized by their large cell size with rabbit α-Repo antibody (red). **B.** mouse α-Cut (red in **B’’**) shows clear signal in the two carpet cells (white arrows). **B’** and **B”.** Carpet cells are recognized by *moody*-Gal4 driving UAS-GFP expression (green). DAPI shows the eye-antennal disc surface. **C.** *Fas2*-Gal4 drives UAS-H2B::RFP (red in **C”**) expression in the two carpet cells (white arrows). **C’** and **C”.** Carpet cells are recognized by their large cell size with DAPI and their location on the posterior edge of the retinal field between the outgoing axons visualized with Phalloidin staining (green). **D.** *Sty*-Gal4 drives UAS-H2B::RFP (red in **D”**) expression in the two carpet cells (white arrows), as well as in other cells in the disc. Due to folding of the imaged disc, the right (**D-I**) and left (**D-II**) carpet cells where not found in the same focal plane. **D’** and **D”**. Carpet cells are recognized by their large cell size with DAPI and their location on the posterior edge of the retinal field between the outgoing axons visualized with Phalloidin staining (green).

In summary, we showed that 5 of the 13 computationally predicted Hb target genes that we tested, were expressed in carpet cells, suggesting that our bioinformatic pipeline allows the identification of new potential regulators of carpet cell development.

## Discussion

### Expression dynamics and clustering recapitulates developmental processes

Although compound eye development is one of the most extensively studied processes in *D. melanogaster*, a comprehensive understanding of genome wide gene expression dynamics is still missing. We performed a genome wide expression study of eye-antennal discs from three larval stages representing late patterning processes and the onset of differentiation (late L2, 72h AEL), differentiation progression (mid L3, 96h AEL) and the completion of the differentiation wave (wandering L3, 120h AEL).

Our data showed that 9,194 of all annotated *D. melanogaster* genes are expressed in the developing eye-antennal disc. We found extensive remodeling of the transcriptomic landscape with 60% of all expressed genes significantly changing their expression profile during the transition from late L2 stages to mid L3 stages. It has been shown that early eye-antennal disc stages are mainly characterized by patterning processes that are necessary to subdivide the initially uniform disc into the organ anlagen for the antennae, the maxillary palps, the compound eyes, the dorsal ocelli and the head cuticle (V. K. L. Merrill, Turner, and Kaufman 1987; Pichaud and Casares 2000; Baonza and Freeman 2002; Aguilar-Hidalgo et al. 2013; Cho et al. 2000; Lebreton et al. 2008; María Domínguez and Casares 2005; Kenyon et al. 2003). Within organ-specific domains, further patterning processes define for instance the dorsal ventral axis in retinal field (Cavodeassi et al. 1999; Yang, Simon, and McNeill 1999; Oros et al. 2010) or the proximal-distal axis of the antennae (Morata 2001). Additionally, the discs grow extensively throughout L1 and L2 stages mainly by cell proliferation (J P Kumar and Moses 2001; Kenyon et al. 2003). With our data, we provide a first glimpse of the gene expression dynamics underlying this fundamental change from predominantly patterning and proliferation processes to the onset of differentiation. Accordingly, the genes active at the late L2 stage were mostly involved in metabolic processes and generation of energy. At the end of L2 stages, the patterning processes are mostly concluded and differentiation starts within each compartment. For instance, in the retinal field the progression of the differentiation wave is accompanied by a reduction in cell proliferation (Wolff and Ready 1991; Jessica E. Treisman 2013). Therefore, mostly genes related to cell differentiation, nervous system development, pattern specification and compound eye development were significantly up- regulated at the mid L3 stage.

On the level of transcriptome dynamics, the transition from the mid L3 stage to late L3 was less pronounced, since only 22% of the expressed genes changed their expression. Interestingly, in this transition again genes related to metabolism and energy production were down-regulated. This can be explained by the fact that at 96h AEL the disc has not yet reached its final size, and cells anterior to the morphogenetic furrow still proliferate (Jessica E. Treisman 2013). Also, directly behind the morphogenetic furrow one last synchronous cell division takes place to give rise to the last cells of the photoreceptor clusters (R1, R6 and R7) (Baonza and Freeman 2002; Jessica E. Treisman 2013). In the light of an ongoing differentiation, the GO terms of genes active at the late L3 stage were also similar to those enriched in the transition from late L2 to mid L3. However, in this case some terms related to later processes were obtained such as R7 cell differentiation or pigment metabolic process, processes taking place late during eye-antennal disc development (Jessica E. Treisman 2013).

The discrepancy between the number of differentially expressed genes in the two studied transitions may in part also be because only female discs were analyzed between 96h and 120h AEL, while we compared mixed males and females at 72h AEL with only females at 96h AEL during the first transition. Since about one third of all genes in *D. melanogaster* show signs of sex-specific expression (Daines et al. 2011), the differentially expressed genes in the first transition may also include some male or female biased genes. This dataset could be an excellent starting point for a comprehensive genome wide analysis of sex-specific gene expression during head development because it has been shown that a strong sexual dimorphism in eye size and head shape exists in *D. melanogaster* (Posnien et al. 2012).

Our clustering of expressed genes based on their dynamic expression profiles resulted in 13 nonredundant clusters (Figure 1), which represent a much more defined representation of the dynamic expression changes during eye-antennal disc development. For example, cluster 7 grouped genes that were similarly high expressed at 72h and 96h AEL, and their expression decreases at 120h AEL. The known genes in this cluster have been described to be related to DNA replication and cell cycle control (Table S2), which corresponds with the fact that active proliferation takes place at these stages (Baonza and Freeman 2002). Thus, other genes that were grouped in this cluster, but for which no previous knowledge is available, are likely also related to these biological functions. Similarly, genes up-regulated in the later stages were separated in more specific clusters, and most of the enriched GO terms are related to differentiation and neuron and eye development. Members of well-known developmental signaling pathways such as EGFR, Notch and cell cycle related genes (e.g. *CycE*) were present in cluster 9 that grouped genes with similarly high expression at 96h AEL and 120h AEL (Figure 1). Among genes, which steadily increased in expression throughout the three studied stages (cluster 11), we found for instance *Delta* (*Dl*), which is one of the Notch receptor ligands (Nicholas E. Baker 2000) and has been shown to fulfill different roles during eye development (Frankfort and Mardon 2002; Kurata et al. 2000; J P Kumar and Moses 2001). Also, *anterior open* (*aop*) (also known as *yan*), which is described to repress photoreceptor differentiation (O’Neill et al. 1994) and also to determine R3 photoreceptor identity (Weber et al. 2008) was present in this cluster. Cluster 4 grouped genes that were highly expressed only at late L3 stage, and correspondingly showed enrichment for genes involved in pigmentation and pupariation (Table S2).

Although, we dissected and sequenced full eye-antennal discs and this tissue contributes to the formation of various organs, the GO enrichment analysis predominantly revealed terms related to general cellular and metabolic processes and retina development. The lack of terms related to antennae or maxillary palp development may be a result of much more extensive research on eye specific developmental processes in comparison to the other organs that develop from the same imaginal disc. However, we revealed various clusters (e.g. clusters 9, 10 and 12) in which GO terms related to leg formation and proximal-distal pattern formation are highly enriched (Table S2). Since antennae and maxillary palps are serially homologue to thoracic appendages, pathways involved in leg, antenna and maxillary palp development are likely to share key regulators (Abu-Shaar and Mann 1998; Campbell and Tomlinson 1998; Dey et al. 2009; Cummins et al. 2003; P. D. S. Dong, Dicks, and Panganiban 2002; Jockusch and Smith 2015; Morata 2001; P. D. Dong, Chu, and Panganiban 2000), suggesting that genes found in these clusters may also play a role in antenna or maxillary palp development.

In summary, we could show that the clustering of genes based on their expression profiles throughout different stages recapitulated the processes underlying eye-antennal disc development exceptionally well. All these observations demonstrate that this comprehensive dataset can be a useful resource to identify new genes involved in the regulation of individual organ development from a common imaginal tissue.

### Identification of central transcription factors involved in eye-antennal disc development

It has previously been shown that co-expression of genes is likely to be a result of regulation by similar or even the same transcription factors (Ideker et al. 2001; Tavazoie et al. 1999; Lee et al. 2002; Allocco, Kohane, and Butte 2004; Altman and Raychaudhuri 2001). This basic assumption has been successfully used to identify central transcriptional regulators in developmental gene regulatory networks (MacArthur et al. 2009; Kemmeren et al. 2014; Deplancke et al. 2006; Ciofani et al. 2012; Yosef et al. 2013; Potier et al. 2014; Junion et al. 2012). The combination of our clustering of dynamic expression profiles with potential transcription factor enrichment within each cluster thus has the potential to reveal key regulators of eye-antennal disc development.

Central and pleiotropic transcriptional regulators are expected to regulate target genes present in different clusters. Accordingly, we found the CREB-binding protein (CBP), also known as Nejire, enriched to regulate genes in nearly all clusters (Figure 1). This zinc-finger DNA binding protein is a co-activator that can act as bridge for other transcription factors to bind specific enhancer elements (Dai et al. 1996; Kwok et al. 1994; McManus and Hendzel 2001), which can explain why we find it to regulate such many target genes. Nejire/CBP has been shown to be involved in many processes during eye development and patterning in *D. melanogaster* (Anderson, Bhandari, and Kumar 2005; Justin P Kumar et al. 2004) and mutations in this gene cause the Rubinstein-Taybi syndrome in humans (Petrif et al. 1995) that among others is characterized by extensive problems during retinal development (van Genderen et al. 2000).

Similarly, the GATA transcription factor Pannier is involved in the establishment of the dorsal-ventral axis of the retinal field of the discs during early L1 and L2 stages (Singh et al. 2005; Singh and Choi 2003), while later during L2 and L3 stages it is known to have a role in defining the head cuticle domain by repressing eye determination genes (Oros et al. 2010; Singh and Choi 2003). In both cases, Pannier is found in a very upstream position of the respective gene regulatory networks that define these cell fates (Maurel-Zaffran and Treisman 2000; Oros et al. 2010). However, despite this well-characterized function during head and eye development little is known about the target genes of Pannier. According to its important central role, we found Pannier enriched to regulate genes in seven out of thirteen clusters. The high number of target genes identified here are prime candidates for further functional analyses to characterize the gene regulatory network downstream of Pannier in more detail.

Besides the very central and general transcriptional regulators, we also identified one very specific cluster that is enriched for genes predominantly regulated by the Ecdysone receptor (EcR) (cluster 5, Figure 1). The fact that this cluster contains mainly genes active at 72h AEL and 120h AEL stages, which represent major stage transitions, confirms that dynamic expression profiling and subsequent clustering can yield highly process specific results. Interestingly, the interpretation of ecdysone related hormonal control has been shown to regulate various aspects of eye-antennal disc development. First, a very general role of ecdysone is to trigger stage transitions, which are characterized by ecdysone hormone pulses before larval molting and pupation (T. Li & Bender, 2000). Second, ecdysone signaling has been shown to promote tissue growth in imaginal discs in general and in the eye-antennal disc specifically (Herboso et al. 2015). Third, the progression of the morphogenetic furrow during eye development is dependent on ecdysone signaling (Brennan et al. 1998). Although the Ecdysone receptor is expressed in the region of the MF, it has later been reported that the ecdysone response is transmitted by the Broad-complex (Brennan et al. 2001; Brennan et al. 1998). Our data provides a set of 282 potential target genes of the Ecdysone receptor and thus represents an excellent starting point to further study the role of ecdysone signaling during eye-antennal disc development. For instance, the target genes could be used to reveal tissue specific genes to understand how a global signal, such as ecdysone can trigger a tissue specific response. Furthermore, our data may be helpful in elucidating the role of the Ecdysone receptor during eye development in *D. melanogaster*.

### Identification of potential novel regulators of eye-antennal disc development

The identification of transcription factors with already well-described central roles during eye-antennal disc development suggests that also new important transcriptional regulators can be identified. For instance, the transcription factor Caudal was found to putatively regulate many genes in the first two clusters of very early expressed genes (Figure 1, cluster 1 and 2). It has been described that Caudal is a downstream core promoter activator (Juven-Gershon, Hsu, and Kadonaga 2008) and very recently it has been found that it cooperates with Nejire to promote the expression of the homeobox gene *fushi tarazu* (*ftz*) (Shir-Shapira et al. 2015). Since Ftz was also found enriched to regulate genes in a cluster of early expressed genes (cluster 1), our results suggest that these three factors could also be acting together during early *D. melanogaster* eye-antennal disc development. Additionally, a Caudal-like transcription factor binding motif has been identified within Sine oculis (So) bound DNA fragments as identified by ChIP-seq (Jusiak et al. 2014), suggesting that So and Cad may co-regulate potential target genes in the eye-antennal disc.

Another unexpected result was the identification of the MADS-box transcription factor Myocyte enhancer factor 2 (DMef2) as being predicted to regulate a number of genes in various clusters. DMef2 is crucial for the development of muscle and heart tissue (Gunthorpe, Beatty, and Taylor 1999). It is expressed in all mesodermal cells during blastoderm stages and its expression gets restricted by the action of the transcription factors Twist and Tinman (Lilly et al. 1994; Nguyen et al. 1994). The detection of *Dmef2* expression in lose cells attached to the developing eye-antennal discs (Figure S2) confirmed that our result is not an artefact, but rather specific. Although these cells are not considered being part of the disc proper, but rather belong to the peripodial membrane, these cells could be precursors of future head muscles. However, some recent findings could hint towards an important role of this transcription factor in eye development. It has recently been reported that DMef2 is implicated in circadian behavior, as it is necessary for the proper fasciculation-defasciculating cycle of neurons (Sivachenko et al. 2013) through one of its target genes *Fasciclin 2* (*Fas2*), which is expressed in some photoreceptor neurons (Mao and Freeman 2009). Additionally, a recent transcriptomics study of larval eye and adult ocelli found that *Dmef2* is expressed in the photoreceptors of both eye types, although the authors did not investigate this finding further (Mishra et al. 2016). These findings certainly encourage additional research on the possible role of DMef2 in photoreceptor cell development.

Taken together, the combination of dynamic gene expression clustering and upstream factor enrichment provides an excellent basis for the identification of potential new regulators involved in a given biological process. Intriguingly, our approach seems to be successful, although the ChIP-seq experiments that identified the direct interaction of a transcription factor with its target genes were not specifically performed in eye-antennal disc tissue at the stages we studied here. Indeed, most data available in current databases is based on experiments in embryonic or adult stages (Celniker et al. 2009; Roy et al. 2010; Nègre et al. 2011; X. Li et al. 2008; Junion et al. 2012). Interestingly, the enrichment of Caudal in clusters 1 and 2 (Table S3) is based on data from a ChIP-seq experiment performed in adult flies (Celniker et al. 2009), but does not represent an experiment performed in embryos (X. Li et al. 2008). This could indicate that Caudal has very different downstream targets during embryogenesis compared to its target genes at later stages. Although this observation may also indicate that the parameters and thresholds used in the different ChIP-seq experiments are very different, a large degree of tissue and stage specific target genes is expected. In the light of this specificity, we may miss eye-antennal disc specific target genes in our survey, but we are confident that one can identify a representative set of target genes to justify further tissue and stage specific ChIP-seq experiments if necessary.

### A new role of Hb in retinal glia development and blood-brain barrier formation

The comprehensive analysis of developmental high-throughput gene expression data in combination with the identification of key upstream regulators suggested that Hb may play an important role during eye-antennal disc development. Using immunostaining and reporter gene expression we confirmed that *hb* is indeed expressed in two large cells in the posterior margin of the eye-antennal discs (Figure 2 and S3). Further co-expression analyses with the pan-glia cell marker Repo and the G-protein coupled receptor Moody indicated that these cells are retinal subperineurial glia cells known as carpet cells (Silies et al. 2007; Bainton et al. 2005) (Figure 3). There are only two carpet cells in each eye-antennal disc and like other subperineurial glia cells they are polyploid and are characterized by huge cell bodies, each spanning half of the retinal field of the eye-antennal discs (Unhavaithaya and Orr-Weaver 2012; Zielke, Edgar, and DePamphilis 2013). It is also known that carpet cells express Moody, a transmembrane protein that is involved in the regulation of the actin cytoskeleton in surface glia and thus influences the positioning of septate junctions (Schwabe et al. 2005). Although the function and some key cellular characteristics of carpet cells are well-understood, their developmental origin and the molecular specification are largely unknown. Our preliminary functional analysis of Hb in carpet cell development and function provided first insights into these open questions.

Upon loss of Hb function (using either RNAi knockdown or Hb^TS^ mutant analysis), the most obvious phenotype was the lack of polyploid cell nuclei in the eye-antennal discs (Figure 5 and S5), indicating that *hb* expression is necessary for the proper development of these cells. Additionally, we showed that the extension of *moody* positive membranes into the eye-antennal discs was impaired in loss of Hb function flies (Figure 6). The carpet cells function as a scaffold for undifferentiated retinal perineurial glia cells, which migrate into the disc to find the nascent axons of differentiating photoreceptors (Silies et al. 2007). In accordance with this scaffold function, we observed regions in the retinal field that were free of perineurial glia cells in eye-antennal discs in which polyploid carpet cell nuclei were not present. This was accompanied by the presence of unorganized axon bundles that did not project properly into the optic stalk (Figure S6). A possible explanation for the patches lacking perineurial glia cells could be the absence of the carpet cell surface to work as support layer for perineurial glia cells. Indeed, we could show that the loss of polyploid carpet cell nuclei was accompanied by impaired formation of *moody* positive cell membranes (Figure 6).

It has been described that in the absence of glia cells, projecting axons are not able to enter the optic stalk or get directed to it (R Rangarajan, Gong, and Gaul 1999). Interestingly, our Hb target gene analysis revealed many candidate target genes with GO terms related to axon guidance (Table S5). A link between undifferentiated retinal glia cells and axon guidance has been established as well. When perineurial glia cells contact newly forming photoreceptor axons, they differentiate into wrapping glia cells and then they enwrap the axons to participate in their projection to the brain lobes (Hummel et al. 2002). Hence, we could not only show impaired carpet cell development upon loss of Hb function, but also observed an impact on glia cell migration and axon guidance as secondary effects.

In contrast to our results, previous studies have shown that carpet cell ablation or a reduction of their size causes over migration of perineurial glia cells anterior to the morphogenetic furrow (Yuva-Aydemir, Bauke, and Klämbt 2011; Silies et al. 2007). The corresponding experiments are based on the induction of cell death in *moody* expressing cells (Silies et al. 2007) and thus affect not only the carpet cells, but also for instance all other subperineurial glia of the brain. Since *hb* expression is very likely specific to carpet cells (see also below), the phenotype obtained here may be more specific. It remains to be studied, however, how carpet cells and subperineurial glia cells of the brain may interact to regulate perineurial glia cell migration in the eye-antennal discs. In many cases, only one carpet cell could be observed in the eye-antennal disc, and this often had a larger polyploid nucleus that was located in the midline of the eye field. In these cases, also no perineurial glia cell over migration could be observed, which might indicate that a single carpet cell could compensate the function of the other missing one.

Since subperineurial glia cells of the brain contribute to the blood-brain barrier (Bainton et al. 2005; Schwabe et al. 2005), we also tested if the loss of Hb function may interfere with blood-brain barrier formation. Indeed, the loss of *hb* expression affected the integrity of the blood-eye barrier, a subset of the blood-brain barrier (Figure 7). This phenotype was not as striking as previously published for *moody* mutant flies (Bainton et al. 2005), where all surface glia cells were affected, suggesting that the carpet cells may indeed only contribute to the retinal part of the blood-brain barrier (i.e. the blood-eye barrier). Intriguingly, the largest portion of the blood-brain barrier is already established by the end of embryogenesis (Beckervordersandforth et al. 2008; von Hilchen et al. 2013), while the eye-antennal disc and developing photoreceptors seem to be accessible for the hemolymph during larval and very early pupal stages. Indeed, the final closure of the blood-brain barrier in the region where the optic stalk contacts the brain (i.e. the blood-eye barrier) is only established late during pupal development (Carlson, Hilgers, and Garment 1998; Carlson et al. 2000). The rather late formation of the blood-eye barrier may be related to the dual role of carpet cells, which first migrate into the eye-antennal disc and only during pupal stages migrate back into the optic stalk towards the brain lobes. By mid-pupa stages they are located at the base of the brain lamina (T. N. Edwards et al. 2012), where they remain throughout adult life and form septate junctions that isolate the brain and retina from the hemolymph (Carlson et al. 2000).

Due to its pivotal role in maintaining the correct physiological conditions in the central nervous system, the blood-brain barrier is of foremost importance for all metazoan organisms. Also in vertebrates, glia cells and especially astrocyte glia, with similar cellular features as subperineurial glia in insects, are the main components of this barrier (Iadecola and Nedergaard 2007). Thus, the study of the function of subperineurial glia cells in blood-brain barrier formation in the invertebrate model *D. melanogaster* can be of great interest to gain insights into central nervous system physiology and disease studies (DeSalvo et al. 2014). Additionally, while the role of Hb in anterior-posterior patterning seems to be conserved only in insects or arthropods (Pinnell 2006; Schröder 2003), its role in central nervous system development is conserved at least across all protostomes (Pinnell 2006). One of the *hb* homologs known in mammals, *ikaros*, which also promotes early-born neuronal fate in mouse (Alsio et al. 2013), has been shown to have a role in conferring identity to retinal progenitor cells (Elliott et al. 2008). It is therefore tempting to investigate a potential role of Ikaros in vertebrate blood-brain barrier development.

### The molecular role of Hb during carpet cell development

The lack of ployploid large carpet cells during larval stages and the loss of blood-brain barrier integrity could either indicate a central role of Hb in specifying carpet cell identity entirely or a more specific role in defining aspects of carpet cell identity such as polyploidy and/or its migratory behavior. The exact role of Hb, however, is still unclear and will require further in-depth analyses. Based on our data presented here, we propose the following cellular functions:

First, the lack of polyploid nuclei could hint towards a role of Hb in regulating the extensive endoreplication process necessary to generate such huge cell nuclei. Indeed, in our target gene analysis we found *(archipelago) ago* as one potential target. Ago has been shown to induce degradation of CyclinE (CycE) (Moberg et al. 2001), a crucial prerequisite for efficient endoreplication cycles (Shcherbata 2004). A role of Hb in the regulation of endoreplication is further supported by a preliminary overexpression experiment. We expressed *hb* ectopically in perineurial glia cells using the specific driver c527-Gal4 (Ito, Urban, and Technau 1995) and we observed an increased number of glia cells with large nuclei in the optic stalk (data not shown). Since retinal perineurial glia cells still proliferate and are undifferentiated (Radha Rangarajan, Courvoisier, and Gaul 2001; R Rangarajan, Gong, and Gaul 1999), the ectopic expression of *hb* may have induced endoreplication cycles, reminiscent of carpet cells.

Second, Hb could be involved in establishing the migratory behavior of carpet cells. In the list of putative Hb target genes, we found many genes with GO terms related to cell migration and, some even specifically with the “glia cell migration” term. Additionally, many of the identified Hb target genes are involved in the epidermal growth factor (EGF) pathway. This is a well-conserved pathway that has received a lot of interest due to its various roles in development and cancer (Gao et al. 2011; Yewale et al. 2013; S. V Sharma et al. 2007). The activation of the EGF receptor (EGFR) by the binding of specific ligands initiates a signaling cascade that transmits information between cells during many different processes, including cell division, differentiation, cell survival and migration (B.-Z. Shilo 2005; B. Shilo 2003). Most of these roles of the EGF pathway have also been shown to be involved in *D. melanogaster* eye development (Malartre 2016). The list of Hb target genes up-regulated at 96h and 120h AEL in eye-antennal disc development includes both positive (*rhomboid*, *Star* and *CBP*) and negative regulators *(Fasciclin2* and *sprouty*) of this pathway. *Fas2* and *sprouty* are specifically expressed in carpet cells (Figure 9C and 9D), suggesting that Hb may actively influence the migratory behavior of carpet cells by activating genes involved in EGFR signaling regulation. Another putative target gene of Hb is *Ets98b*.

Intriguingly, it has recently been shown during early embryonic development in the common house spider *Parasteatoda tepidariorum* that the ortholog Ets4/Ets98b induces ectopic cell migration upon misexpression (Pechmann et al., submitted). We also performed preliminary misexpression experiments and expressed *hb* ectopically in wrapping glia cells (not shown). In such misexpression eye-antennal discs we observed cell nuclei between the axon bundles in the optic stalk. These may be wrapping glia cells that over migrate into the stalk, although they normally remain in the eye-antennal disc and only their extended cell membranes project to the brain lamina or medulla to accompany the photoreceptor axons (Hummel et al. 2002).

In summary, our functional analyses in combination with computationally supported target gene prediction suggests that Hb plays a central role in specifying key cellular features of carpet cells: polyploidy and extensive migratory abilities.

### Implication on the origin and nature of carpet cells

Although carpet cells fulfill fundamental functions, it is still unclear where these cells originate from. Based on observations by Choi and Benzer (1994) using the enhancer trap line M1-126, these cells originate in the optic stalk where they are present at late L2 stage (Choi and Benzer 1994). It has also been proposed that carpet cells may originate from a pool of neuroblasts in the neuroectoderm during embryogenesis (Homem and Knoblich 2012) or in the optic lobes (Apitz and Salecker 2014). A clonal analysis using the FLP-out system suggests that various retinal glia cell types, including the carpet cells, originate from at least one mother cell at L1 larval stage (R Rangarajan, Gong, and Gaul 1999). Since only one polyploid cell nucleus seems to originate from one clone (R Rangarajan, Gong, and Gaul 1999) and we show that in some loss of Hb imaginal discs only one polyploid cell nucleus is present, we propose that the two carpet cells may originate independently probably from two mother cells defined during L1 stages. Hb may be the key transcription factor specifying carpet cell fate to distinguish them from other retinal glia cells. Our observation that loss of Hb function resulted in loss of polyploid carpet cell nuclei when Hb^TS^ mutant flies were transferred to the restrictive temperature during the L1 larval stage, further supports an involvement of Hb during this stage (Figure 5 and S5)

Carpet cells have been shown to be a sub-population of the subperineurial glia cells due to shared key cellular features, such as the formation of extensive septate junctions (Silies et al. 2007). However, a well-established subperineurial glia cell driver (NP2276 (Awasaki et al. 2008)) does drive reporter gene expression in brain subperineurial glia, but not in carpet cells (data not shown). In contrast, we only detected *hb* expression in carpet cells and not in any subperineurial glia cells in the larval brain (results confirmed both using immunostaining and two driver lines (VT038544 and VT038545), not shown). Additionally, if we compare our list of putative Hb target genes with the 50 genes enriched in adult blood-brain barrier surface glia (DeSalvo et al. 2014), we only find *Fas2* to be present in both datasets. All these data suggest that carpet cells are indeed a retina specific subperineurial glia cell type that is molecularly very distinct from brain subperineurial glia cells.

The use of the newly analyzed driver lines VT038544 and VT038545, which drive expression specifically in the carpet cells in combination with the extensive list of potential Hb target genes, of which many are likely to be expressed in this specific glia cell type, represents a valuable resource to address the questions concerning the origin of these cells in more detail.

### Conclusions

In this study, we identified a new role of Hb in retinal glia cell development. This finding has only been possible because we studied the dynamic expression profiles of all genes expressed during eye-antennal disc development. Since the RNA levels of *hb* in the entire eye-antennal disc are negligible, we could identify Hb as central factor only through the expression profiles of its putative target genes, which are steadily up regulated throughout development. This up regulation is very likely due to the large cell bodies of the carpet cells, which need to produce high amounts cytosolic or membrane bound proteins. At earlier stages, carpet cells are not yet in the eye-antennal discs, and *hb* expression could have only been identified by studies focused on the optic stalk. Moreover, we could show that refining the putative Hb target genes by incorporating the expression data results in a list that contains genes with GO terms highly specific for the putative function of Hb in carpet cells. Based this stepwise identification of target genes, we could select and confirm a high number of those experimentally.

All these findings demonstrate that the combination of high throughput transcript sequencing with a ChIP-seq data based transcription factor enrichment analysis can reveal previously unknown factors and also their target genes, and therefore increase the number of connections within the underlying developmental GRNs. Other studies have searched for regulating transcription factors that were in the same co-expression clusters as its targets genes (Potier et al. 2014). However, upstream orchestrators do not necessarily have the same expression levels as their targets. Therefore, the combination of ChIP-seq methods in RNA-seq co-expression analyses has proven to be a powerful tool to identify new developmental regulators that can complement other studies based on reverse genetics.

## Materials and methods

### RNA extraction and sequencing

*D. melanogaster* (OregonR) flies were raised at 25°C and 12h:12h dark:light cycle for at least two generations and their eggs were collected in 1h windows. Freshly hatched L1 larvae were transferred into fresh vials in density-controlled conditions (30 freshly hatched L1 larvae per vial). At the required time point, eye-antennal discs of either only female larvae (96h and 120h AEL) or male and female larvae (72h AEL) were dissected and stored in RNALater (Qiagen, Venlo, Netherlands). 40-50 discs were dissected for the 120h samples, 80-90 discs for the 96h samples and 120-130 discs for the 72h samples. Three biological replicates were generated for each sample type.

Total RNA was isolated using the Trizol (Invitrogen, Thermo Fisher Scientific, Waltham, Massachusetts, USA) method according to the manufacturer’s recommendations and the samples were DNAseI (Sigma, St. Louis, Missouri, USA) treated in order to remove DNA contamination. RNA quality was determined using the Agilent 2100 Bioanalyzer (Agilent Technologies, Santa Clara, CA, USA) microfluidic electrophoresis. Only samples with comparable RNA integrity numbers were selected for sequencing.

Library preparation for RNA-seq was performed using the TruSeq RNA Sample Preparation Kit (Illumina, catalog ID RS-122-2002) starting from 500 ng of total RNA. Accurate quantitation of cDNA libraries was performed using the QuantiFluor™dsDNA System (Promega, Madison, Wisconsin, USA). The size range of final cDNA libraries was determined applying the DNA 1000 chip on the Bioanalyzer 2100 from Agilent (280 bp). cDNA libraries were amplified and sequenced (50 bp single-end reads) using cBot and HiSeq 2000 (Illumina). Sequence images were transformed to bcl files using the software BaseCaller (Illumina). The bcl files were demultiplexed to fastq files with CASAVA (version 1.8.2).

## BIOINFORMATICS ANALYSES

### Quality control

Quality control was carried out using FastQC software (version 0.10.1, Babraham Bioinformatics). All samples but one (“72hC” sample) had quality score >Q28 for all read positions. 12% of reads in sample “72hC” had an “N” in position 45, probably due to an air bubble in the sequencer. Following recently published guidelines (MacManes 2014; Williams et al. 2016), sequences were not trimmed but the aligner software was used to this purpose instead with very stringent parameters (see below). All raw fastq files are available through GEO Series accession number GSE94915.

### Read mapping

Transcript sequences (only CDS) of *D. melanogaster* (r5.55) were downloaded from FlyBase and only the longest transcript per gene was used as reference to map the reads using Bowtie2 (Langmead and Salzberg 2012) with parameters `–very–sensitive–local –N 1`. The number of reads mapping to each transcript were summarized using the command `idxstats` from SAMtools v0.1.19 (H. Li et al. 2009). A summary of raw read counts mapped to each gene and time point is available at the GEO repository (GSE94915).

### Gene expression clustering

HTSFilter (Rau et al. 2013) was used with default parameters to discard lowly expressed genes across all samples. The function `PoisMixClusWrapper` from the library HTSCluster (Rau et al. 2015) was applied on the rest of genes with the parameters: `gmin=1, gmax=25, lib.type=“DESeq”`.

Different model selection approaches are used by HTSCluster (i.e. to identify the number of clusters that best describe the data (see (Rau et al. 2015)). Our previous experience with this package had shown that the BIC and ICL methods report always as many clusters as we have allowed to test for (corresponding to the “gmax” parameter). Also for this analysis, both methods reported 25 as the most likely number of clusters, which was the input “gmax” value. Consequently, we discarded these results and we only analyzed the results of the methods that use slope heuristics to calculate the best number of clusters, namely DDSE and Djump. The DDSE method reported 19 clusters, with 8,626 genes having MAP > 99% while the Djump method reported 13 clusters, with 8,836 genes having MAP > 99%. Careful inspection of the lambda values of each of these clusters showed that the additional clusters predicted by the DDSE method presented negligible variation to the 13 clusters predicted by Djump. Additionally, GO term analysis (see below) of the genes in the 19 clusters predicted by DDSE showed redundant terms for the very similar additional clusters, which was not the case with the 13 clusters predicted by Djump. Therefore, we concluded that the additional clusters present in the DDSE prediction were unlikely to represent significant biological differences and that the 13 clusters predicted by Djump could sufficiently describe the profiles of the groups of co-expressed genes and we used them for all following analyses.

Genes with predicted MAP < 99% were discarded. Cluster assignment results can be found at the GEO repository (GSE94915). For the plots, the variance stabilizing transformation from DESeq2 (Love, Anders, and Huber 2014) library was used to normalize the background read count of the genes belonging to each cluster.

The Gene Ontology terms enriched in each cluster of genes were obtained with the plugin BiNGO (Maere, Heymans, and Kuiper 2005) in Cytoscape v3.1.1 (Cline et al. 2007) with default parameters. The ontology terms and corresponding *D. melanogaster* annotations were downloaded from geneontology.org (Ashburner et al. 2000; Consortium 2015) (as of January 2015).

The transcription factors enriched to regulate the genes of each cluster were obtained with the i-cisTarget method (Herrmann et al. 2012) with the following parameters: dm3 assembly, only “TF binding sites”, 5 Kb upstream and full transcript as mapping region, 0.4 as minimum fraction of overlap, 3.0 as NES threshold and 0.01 ROC threshold.

### Identification of Hb target genes

The i-cisTarget method (Herrmann et al. 2012) to detect transcription factor enrichment in the regulatory regions of co-regulated genes is based on the arbitrary partition of the *D. melanogaster* genome in more than 13,000 regions. All genes included in a particular region are associated to the transcription factor binding interval, resulting maybe in an unspecific association between transcription factor and target genes. Therefore, we aimed to generate a more confident list of putative Hb target genes in the eye-antennal disc. From Berkeley *Drosophila* Transcription Network Project (BDTNP) site (X. Li et al. 2008), BED files were downloaded for the Hb (anti-Hb (antibody 2), stage 9) ChIP-chip experiment (Symmetric-null test and 1% FDR cutoff). The LiftOver tool from UCSC Browser (Kent et al. 2002) was used to transform the dm2 coordinates into the dm3 assembly. The closest gene to each ChIP-chip interval was identified with the script `annotatePeaks.pl` from the HOMER suite of tools (Heinz et al. 2010). Enrichment for the Hb motif in the regulatory regions of the identified genes were confirmed with the script `findMotifGenome.pl` from the same suite. The identified enriched Hb motif (as matrix) was used to search again the closest genes to the ChIP-chip intervals using the script `annotatePeaks.pl` with the parameters tss –size –1000,1000 –m *motif_matrix*. The genes with at least one instance of the motif were selected as Hb high confident targets. Cytoscape v3.1.1 (Cline et al. 2007) was used to visualize high confidence Hb targets which are significantly up- and down-regulated in the 72h AEL to 96h AEL and 96h AEL to 120h AEL transitions.

## EXPERIMENTAL PROCEDURES

### Fly lines and crosses

The following fly lines were used: UAS-*hb*_dsRNA_ (Bloomington Stock Center #54478, #29630 and #34704 and Vienna Drosophila Research Center #107740), *hb*-Gal4 (Vienna Tile library (Pfeiffer et al. 2008) VT038544 and VT038545), UAS-*hb* (Bloomington Stock Center #8503), *repo*-Gal4/TM6B (kindly provided by Marion Sillies), *moody*-Gal4 ((Schwabe et al. 2005) kindly provided by Christian Klämbt), *DMef2*-Gal4 (Bloomington Stock Center #25756) UAS-stinger-GFP (nGFP) ((Barolo, Carver, and Posakony 2000) kindly provided by Gerd Vorbrüggen), UAS-H2B:RFP (kindly provided by Andreas Wodarz) and 20xUAS-mCD8::GFP (Bloomington Stock Center #32194). Lines expressing Gal4 under control of regulatory regions of the Hb putative target genes were obtained from Bloomington Stock Center (*ago*-Gal4 (#103-788), *brk*-Gal4 (#53707), *CadN*-Gal4 (#49660), *DI*-Gal4 (#45495), *Fas2*-Gal4 (#48449), *kni*-Gal4 (#50246), *rho*-Gal4 (#49379), *robo3*-Gal4 (#41256), *Sox21b*-Gal4 (#39803), *Src64B*-Gal4 (#49780), *sty*-Gal4 (#104304) and *tkv*-Gal4 (#112552)).

All crosses were performed with an approximate ratio of 4:3 female:male flies. Crosses were always provided with additional yeast and were kept at 12h:12h dark:light cycle and controlled humidity, except the RNAi experiments, that were kept at 28°C and constant darkness.

### *hb* RNA interference

We obtained 4 different UAS-*hb*_dsRNA_ lines from Bloomington Stock Center (#54478, #29630 and #34704) and from the Vienna Drosophila Research Center (#107740). We took advantage of the fact that Hb is known to be necessary during early embryogenesis (Lehmann and Nüsslein-Volhard, 1987; Nüsslein-Volhard and Wieschaus, 1980) to evaluate the knock-down efficiency. UAS-*hb*_dsRNA_ flies were crossed to the *hb*-Gal4 lines (VT038544 and VT038545) to see if the survival of the offspring was affected. Only one of the RNAi lines, namely #34704, produced no adult flies and few dead pupae when crossed with the *hb*-Gal4 flies. The other three lines produced a normal number of offspring with no obvious phenotype. Consequently, we used the #34704 line for the knock-down experiments. Please note that the evaluation of knock-down efficiency in the developing eye-antennal discs using quantitative PCR is very limited because the expression of *hb* itself is very low (practically no reads are detected by RNA-seq, not shown).

### Hb^TS^ cross

*Hb^TS1^, rsd^1^/TM3, Sb^1^* flies (Bloomington Stock Center #1753) were crossed to *hb^12^, st^1^, e^1^/TM3, Sb^1^* flies (Bloomington Stock Center #1755) to generate a *hb^TS1^/hb^12^* stock. This line was kept at 18°C and constant light and larvae were only transferred to the restrictive temperature (28°C) for the loss of function experiments.

### *in-situ* hybridization

Standard procedures were followed to clone a fragment (872 bp) of *hunchback* gene sequence into pCRII vector and to synthesize an antisense digoxigenin-labeled RNA probe (and sense probe for the negative control). RNA probes were hydrolyzed with Na-Carboante buffer for 30.5 minutes. Eye-antennal discs were dissected in cold PBS and fixed with 4% paraformaldehyde. Hybridization was carried out at 63°C overnight with 5 μl of RNA probe (291 ng/μl) in 50 μl of hybridization buffer. Anti-Dig antibody (1:2000, Sigma-Aldrich) was used to detect the probe and revealed with NBT/BCIP reaction mix. Pictures were taken with a Zeiss Axioplan microscope.

### Immunohistochemistry

Antibody stainings were performed using standard procedures (Klein 2008). In all cases, dissected eye-antennal discs were fixed with 4% paraformaldehyde before incubating with primary and secondary antibodies. Antibodies used were: rabbit α-Repo ((von Hilchen et al. 2013), 1:1000), guinea-pig α-Hb ((Kosman, Small, and Reinitz 1998), 1:50), mouse α-cut (Invitrogen, 1:100), rabbit α-Hb (kind gift from Chris Q. Doe, 1:100), Cy3-α-HRP (kind gift from Martin Göpfert, 1:300), goat α-rabbit Alexa Fluor 488 (Invitrogen, 1:1000), goat α-rabbit Alexa Fluor 555 (Invitrogen, 1:100) and goat α-guinea-pig Alexa Fluor 555 (Invitrogen, 1:1000). A solution of 80% glycerol + 4% n-propyl gallate in PBS was used as mounting medium for all stained discs. Pictures were taken on a Zeiss LSM-510 confocal laser scanning microscope.

### Blood-eye barrier assay

The integrity of the blood-eye barrier of *hunchback* knock-down flies was studied following the protocol from (Pinsonneault et al. 2011). *moody*-Gal4 virgin females were crossed with UAS-*hb*_dsRNA_ males at 28°C. UAS-*hb*_dsRNA_ flies were used as control and also raised at 28°C. 2-3 day old adults from these crosses were injected in the abdomen with 3-5 kDa FITC dextran (Sigma-Aldrich) (0.3 μl the females and 0.2 μl the males of 25 mg/ml solution). Animals were allowed to recover in fresh food over-night. Only surviving animals were scored. Dye penetrance in each eye was assessed qualitatively using a LEICA M205 FA fluorescent stereo microscope.

## Abbreviations

AEL: after egg laying
L1: first larval stage
L2: 2nd larval stage
L3: third larval stage
Hb: Hunchback
MF: morphogenetic furrow

## Acknowledgements

We would like to thank Alistair McGregor, Isabel Almudi, Marion Silies, Iris Salecker, Gerd Vorbrüggen, Fernando Casares, Ernst Wimmer and Halyna Shcherbata for providing reagents and for valuable discussions and critical comments throughout the project. We thank Marita Büscher for various discussions about *Drosophila* nervous system development. We would also like to thank the Transcriptome Analysis Lab (TAL) (University Medical Center Göttingen, UMG) in Göttingen for the Illumina sequencing. Stocks obtained from the Bloomington Drosophila Stock Center (NIH P40OD018537) were used in this study.

This work was funded by a grant of the Volkswagen Foundation (project number: 85 983) and by the Emmy Noether Programme of the Deutsche Forschungsgemeinschaft (grant number: PO 1648/3-1) to NP, and a “la Caixa-DAAD” fellowship to MT-O. The work of MT-O was additionally kindly supported by the Göttingen Graduate School for Neurosciences, Biophysics, and Molecular Biosciences (GGNB).

## Authors contribution

NP and MT-O conceived the experiments. MT-O extracted the RNA for Illumina sequencing and performed all bioinformatics analyses. MT-O, JS, GW and FK performed the functional Hunchback experiments. MT-O and NP interpreted the data and wrote the manuscript. All authors read and approved the manuscript.

---

**Supplementary Figure 1.**
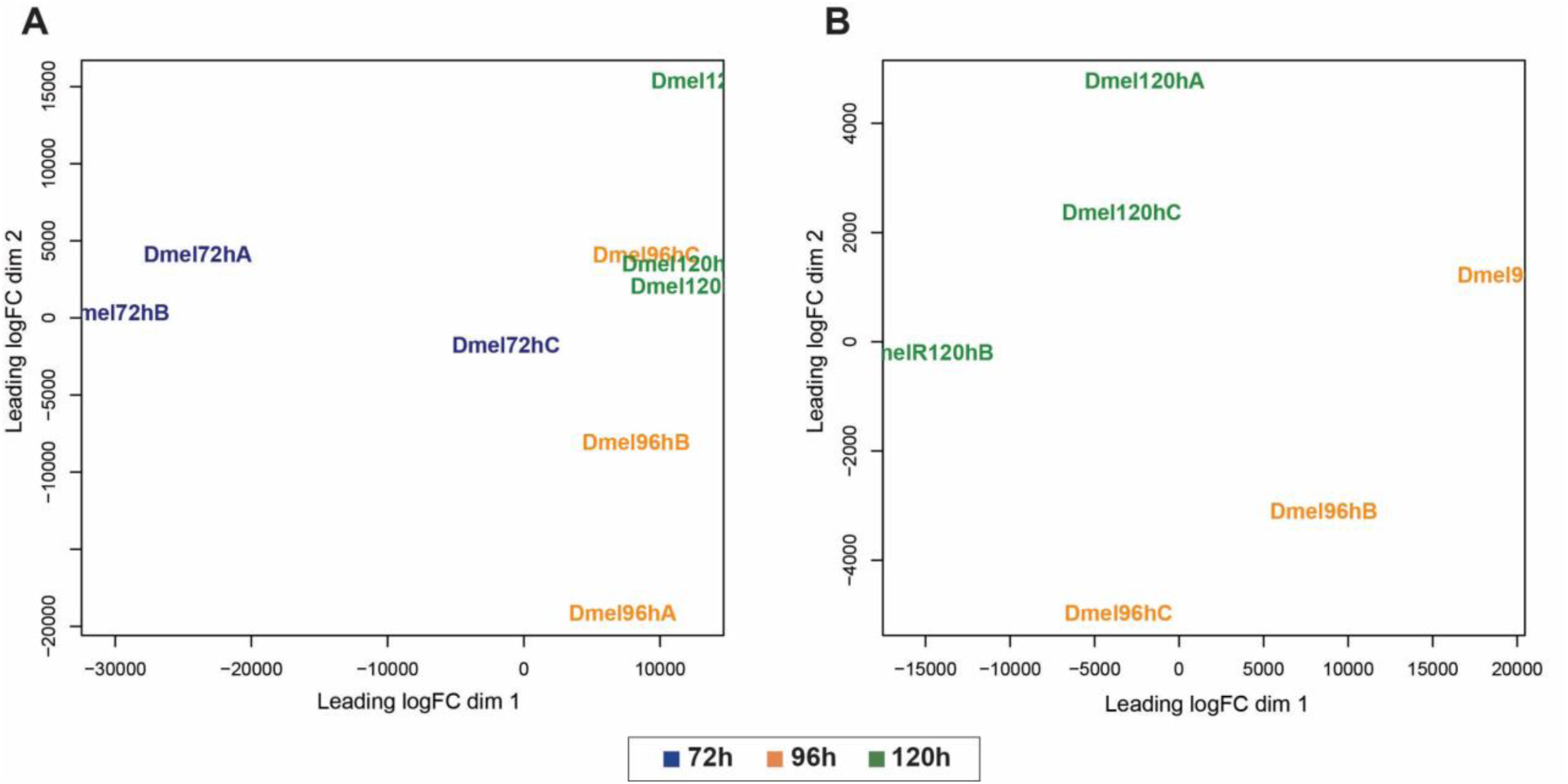
Multi-dimension scaling plot of RNA-seq samples. **A.** Count data of all three time points (72h AEL, 96h AEL and 120h AEL). **B.** Count data of only 96h AEL and 120h AEL.

**Supplementary Figure 2.**
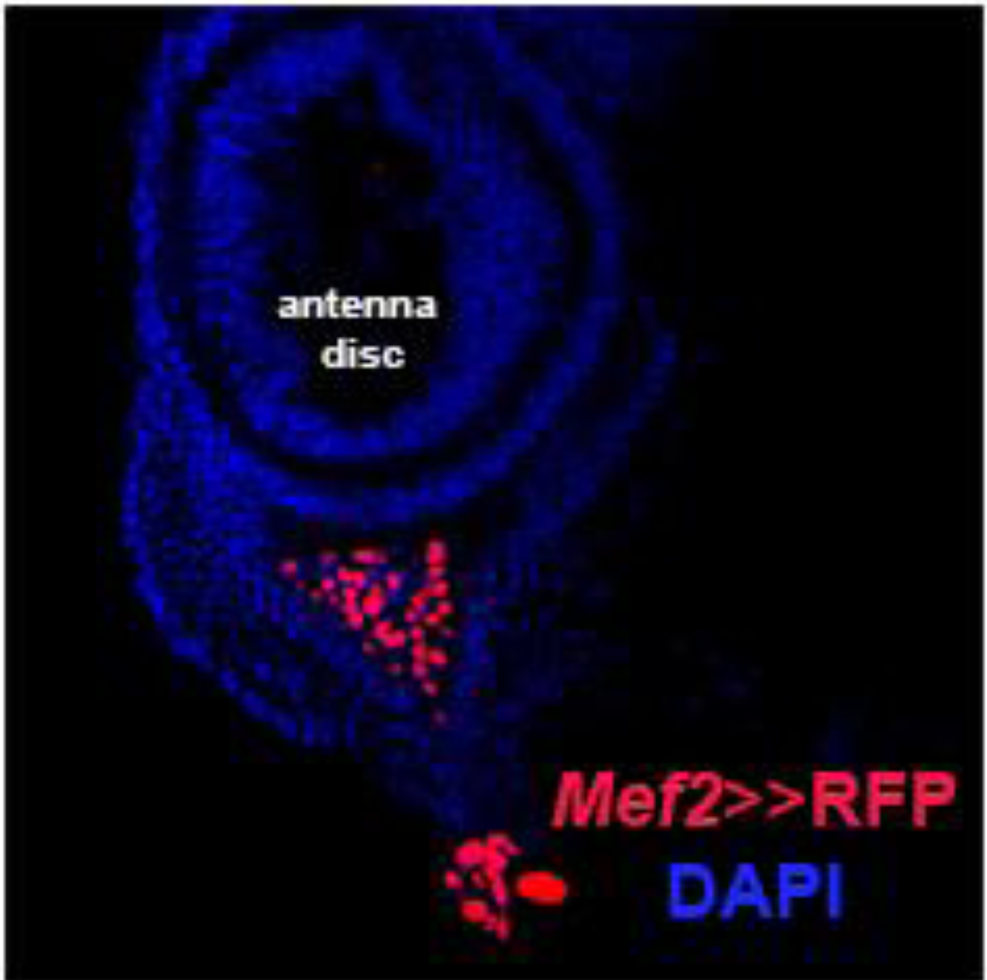
Mef2 driver line expression. *Mef2*-expressing cells are visualized with a *Mef2*-Gal4 driver line crossed with UAS-H2B::RFP reporter (red).

**Supplementary Figure 3.**
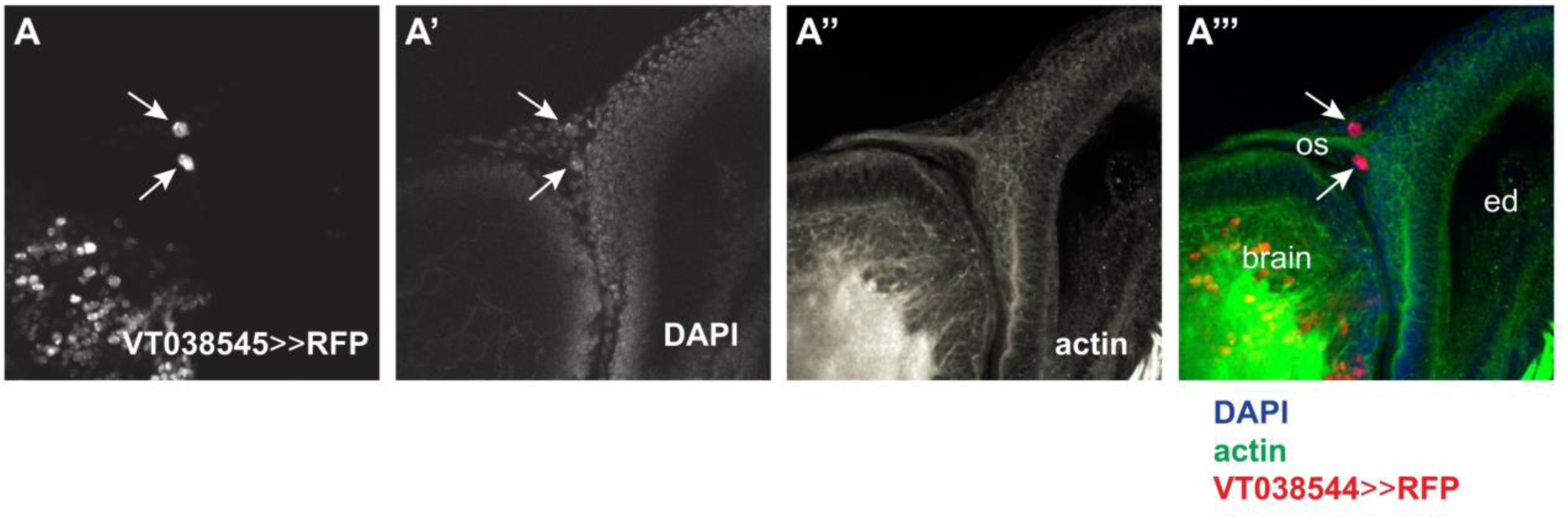
VT038545 (*hb*-Gal4) driver line expression in late L3 eye-antennal discs. Driver line VT038545-Gal4 drives UAS-H2B::RFP expression in the two carpet cells. Anterior is to the right. Eye disc (ed), optic stalk (os).

**Supplementary Figure 4.**
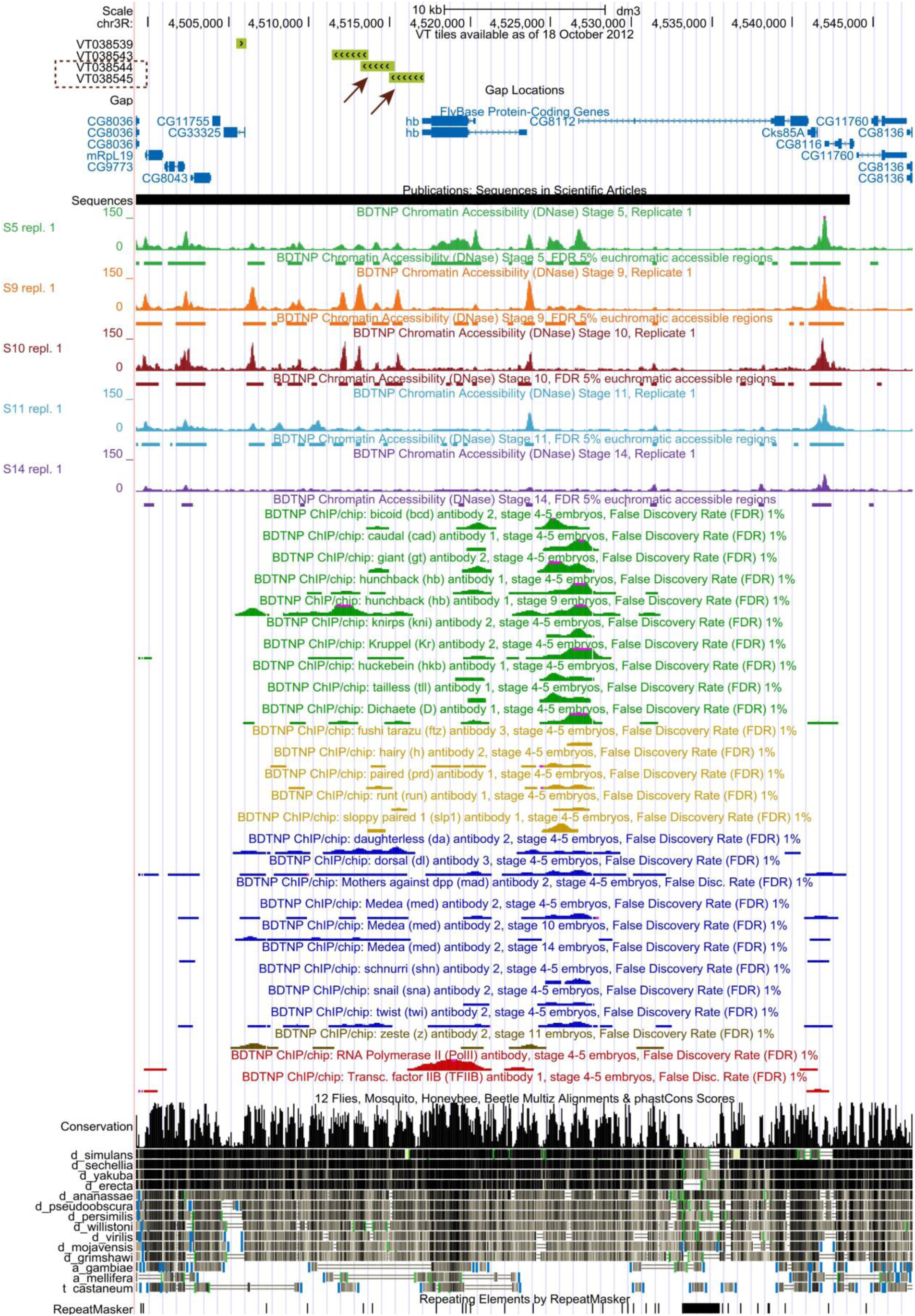
Genomic location of Vienna Tile *hb* driver lines. Arrows indicate the regions used to drive *hb* expression with Gal4 system. Bellow, are colored tracks provided by the BDTNP project (X. Li et al. 2008) showing open chromatin profiles and transcription factor binding. The last black tracks show sequence conservation across different insect species. These tracks were visualized using UCSC Browser (Kent et al. 2002).

**Supplementary Figure 5.**
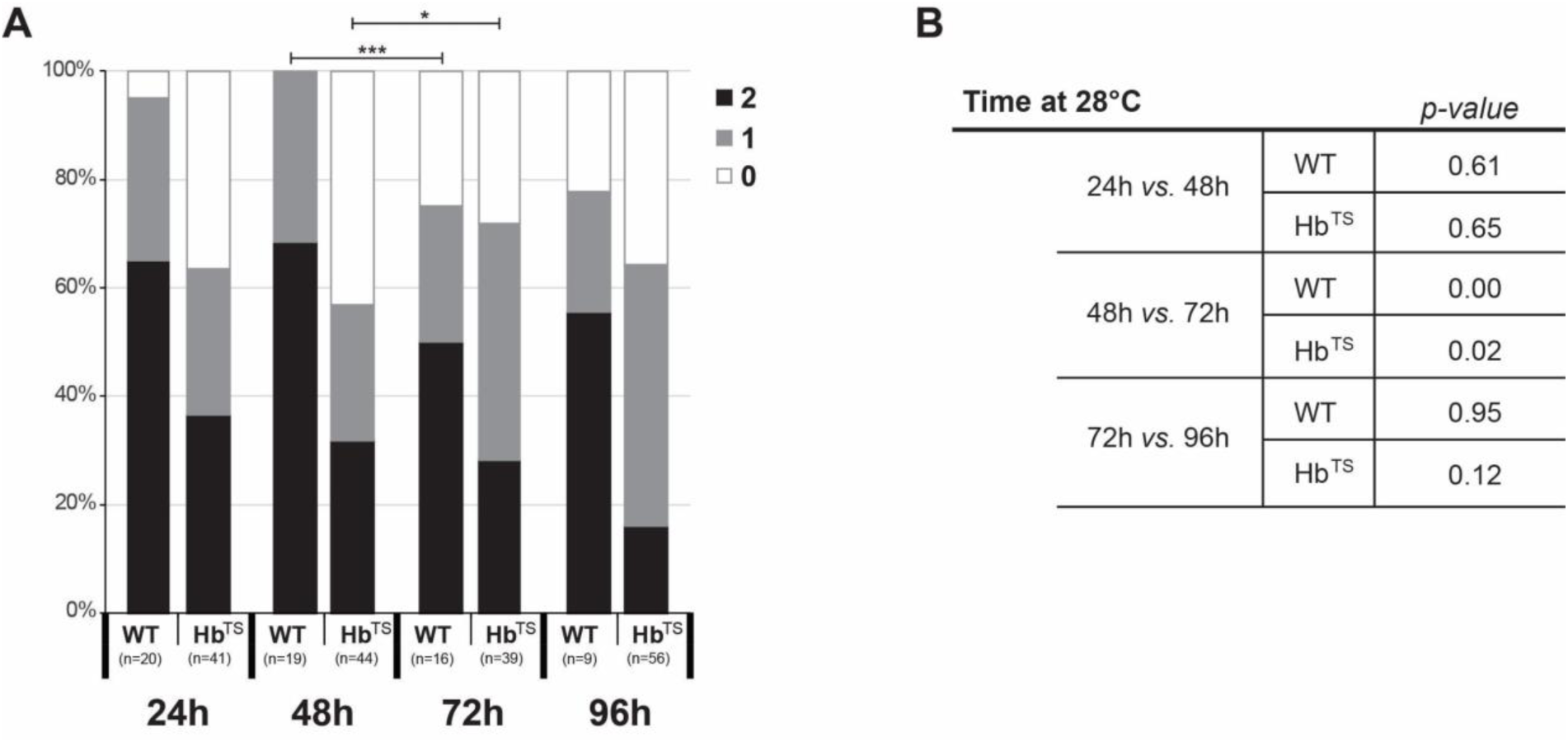
The strength of the effect of loss of Hb function in carpet cells is not significantly different at different time points. **A.** A significant difference in the distribution of the number of polyploid glia cells in Hb^TS^ flies is only observed between raising larvae at the restrictive temperature 48h AEL and 72h AEL. However, this difference is also significant in the wild type (WT). This can be due to the fact that more larvae die when transferred to the restrictive temperature too early (at 24h AEL or 48h AEL). **B.** Pearson’s Chi-squared test was performed to determine if the distribution of the different number of cells (0, 1 or 2) was equal across the time points for the same conditions (WT or HB^TS^). *: p-val < 0.05, ***: p-val < 0.0005.

**Supplementary Figure 6.**
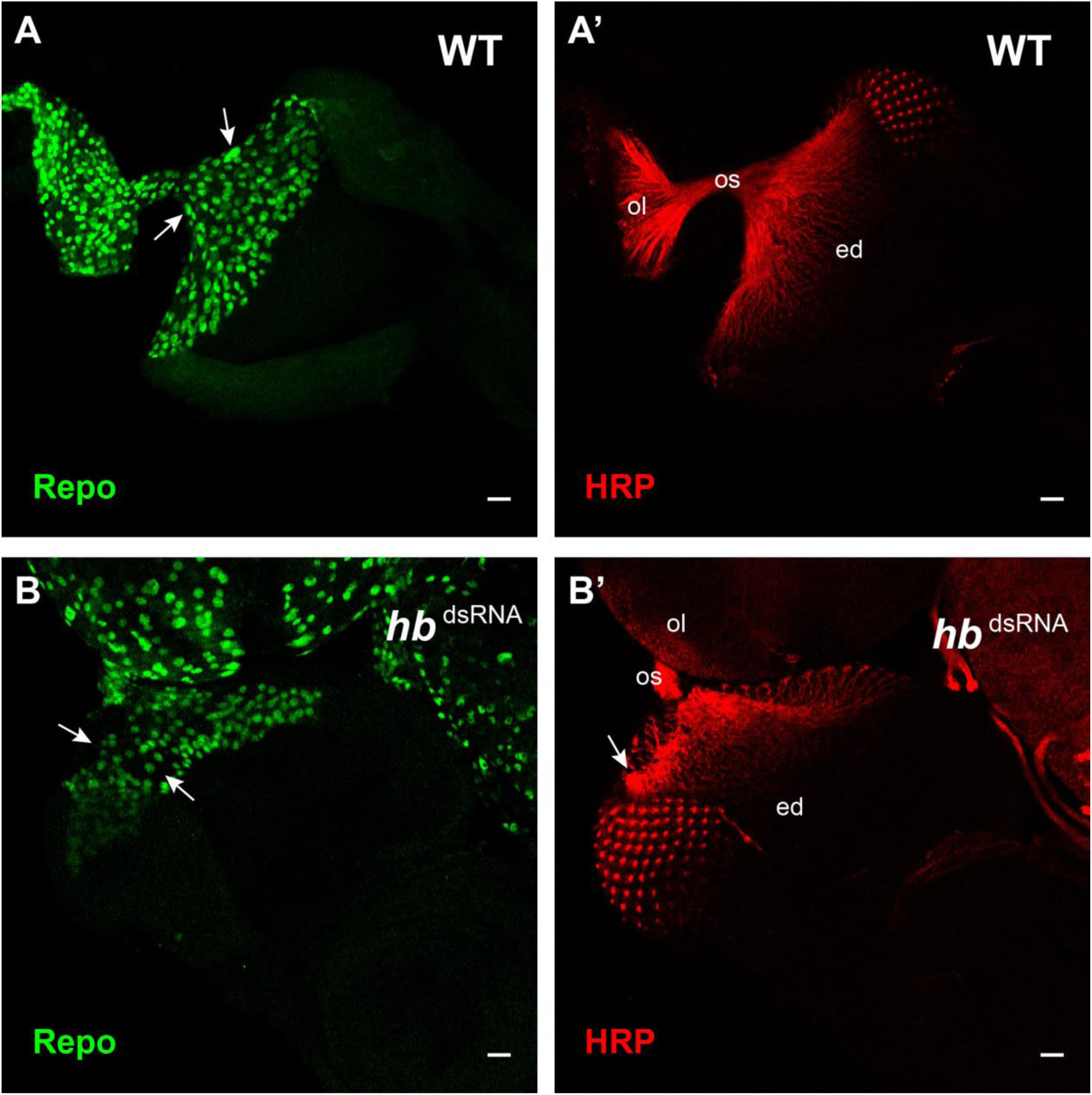
Hb loss of function affects axon projection and the organization of other retinal glia cells. Late L3 eye-antennal discs attached to the optic lobe immunostained with rabbit α-Repo (green) and Cy3-conjugated-HRP (red) antibodies. Eye disc (ed), optic stalk (os), optic lobe (ol). Scale bar = 20 μm. **A**. In wild type larvae, glia cells occupy all the basal surface of the eye-antennal disc posterior to the morphogenetic furrow to support the developing photoreceptors and their axons. Carpet cell nuclei can be observed at the posterior margin of the eye-antennal disc (white arrows). **A’**. Axons project in an organized manner from the developing photoreceptors in the eye-antennal disc into the optic lobes through the optic stalk. **B**. In *repo*>>*hb*^dsRNA^ larvae, patches without glia cells can be observed in the basal surface of the eye-antennal disc (white arrow), and carpet cell nuclei cannot be identified. **B’**. Axons do not project correctly and form unorganized bundles (white arrows).

**Supplementary Table 1. Differentially expressed genes.**

**Supplementary Table 2. Significantly enriched GO terms in the expression clusters.**

**Supplementary Table 3. Significantly enriched transcription factors in the expression clusters.**

**Supplementary Table 4. Putative Hb target genes differentially expressed.**

Table contains two sheets: first sheet lists putative Hb targets differentially expressed between 72h AEL and 96h AEL and second sheet lists the differentially expressed genes between 96h AEL and 120h AEL. “Instances”: number of Hb motifs found ±1000 bp from TTS. Right-side table shows how many of these genes belong to each cluster and the percentage over the total number of genes in that cluster.

**Supplementary Table 5. Putative Hb target genes in clusters 12 and 13.**

Table contains three sheets: first sheet contains the gene ID, name and symbol of the 77 genes, and the cluster they belong to; second sheet lists the GO terms associated to each of the 77 genes; third sheet contains the number of times each GO term appears in the second sheet.

## References

Abu-Shaar, M, and R S Mann. 1998. “Generation of Multiple Antagonistic Domains along the Proximodistal Axis during Drosophila Leg Development.” Development (Cambridge, England) 125 (19): 3821–30.

Aerts, Stein, Xiao Jiang Quan, Annelies Claeys, Marina Naval Sanchez, Phillip Tate, Jiekun Yan, and Bassem a. Hassan. 2010. “Robust Target Gene Discovery through Transcriptome Perturbations and Genome-Wide Enhancer Predictions in Drosophila Uncovers a Regulatory Basis for Sensory Specification.” PLoS Biology 8 (7). doi:10.1371/journal.pbio.1000435.g001.

Aguilar-Hidalgo, Daniel, María a Domínguez-Cejudo, Gabriele Amore, Anette Brockmann, María C Lemos, Antonio Córdoba, and Fernando Casares. 2013. “A Hh-Driven Gene Network Controls Specification, Pattern and Size of the Drosophila Simple Eyes.” Development (Cambridge, England) 140 (1): 82–92. doi:10.1242/dev.082172.

Allocco, Dominic J, Isaac S Kohane, and Atul J Butte. 2004. “Quantifying the Relationship between Co-Expression, Co-Regulation and Gene Function.” BMC Bioinformatics 5 (1): 18. doi:10.1186/1471-2105-5-18.

Alsio, J. M., Basile Tarchini, Michel Cayouette, and Frederick J Livesey. 2013. “Ikaros Promotes Early-Born Neuronal Fates in the Cerebral Cortex.” Proceedings of the National Academy of Sciences 110 (8): E716–25. doi:10.1073/pnas.1215707110.

Altman, R. B., and S. Raychaudhuri. 2001. “Whole-Genome Expression Analysis: Challenges beyond Clustering.” Current Opinion in Structural Biology 11 (3): 340–47. doi:10.1016/S0959-440X(00)00212-8.

Anais Tiberghien, Marie, Gaelle Lebreton, David Cribbs, Corinne Benassayag, and Magali Suzanne. 2015. “The Hox Gene Dfd Controls Organogenesis by Shaping Territorial Border through Regulation of Basal DE-Cadherin Distribution.” Developmental Biology 405 (2). Elsevier: 183–88. doi:10.1016/j.ydbio.2015.07.020.

Anderson, Jason, Rohan Bhandari, and Justin P. Kumar. 2005. “A Genetic Screen Identifies Putative Targets and Binding Partners of CREB-Binding Protein in the Developing Drosophila Eye.” Genetics 171 (4): 1655–72. doi:10.1534/genetics.105.045450.

Anderson, Jason, Claire L. Salzer, and Justin P. Kumar. 2006. “Regulation of the Retinal Determination Gene Dachshund in the Embryonic Head and Developing Eye of Drosophila.” Developmental Biology 297 (2): 536–49. doi:10.1016/j.ydbio.2006.05.004.

Apitz, Holger, and Iris Salecker. 2014. “A Challenge of Numbers and Diversity: Neurogenesis in the Drosophila Optic Lobe.” Journal of Neurogenetics 28 (3–4): 233–49. doi:10.3109/01677063.2014.922558.

Ashburner, Michael, Catherine A. Ball, Judith A. Blake, David Botstein, Heather Butler, J. Michael Cherry, Allan P. Davis, et al. 2000. “Gene Ontology: Tool for the Unification of Biology.” Nature Genetics 25 (1). Nature Publishing Group: 25–29. doi:10.1038/75556.

Atkins, Mardelle, and Graeme Mardon. 2009. “Signaling in the Third Dimension: The Peripodial Epithelium in Eye Disc Development.” Developmental Dynamics 238 (9): 2139–48. doi:10.1002/dvdy.22034.

Awasaki, Takeshi, Sen-Lin Lai, Kei Ito, and Tzumin Lee. 2008. “Organization and Postembryonic Development of Glial Cells in the Adult Central Brain of Drosophila.” The Journal of Neuroscience: The Official Journal of the Society for Neuroscience 28 (51): 13742–53. doi:10.1523/JNEUROSCI.4844-08.2008.

Bainton, Roland J., L. T Y Tsai, Tina Schwabe, Michael DeSalvo, Ulrike Gaul, and Ulrike Heberlein. 2005. “Moody Encodes Two GPCRs That Regulate Cocaine Behaviors and Blood-Brain Barrier Permeability in Drosophila.” Cell 123 (1): 145–56. doi:10.1016/j.cell.2005.07.029.

Baker, N E, and S Y Yu. 2001. “The EGF Receptor Defines Domains of Cell Cycle Progression and Survival to Regulate Cell Number in the Developing Drosophila Eye.” Cell 104 (5): 699–708. doi:10.1016/S0092-8674(02)06076-2.

Baker, Nicholas E. 2000. “Notch Signaling in the Nervous System. Pieces Still Missing from the Puzzle.” BioEssays 22 (3): 264–73. doi:10.1002/(SICI)1521-1878(200003)22:3<264::AID-BIES8>3.0.CO;2-M.

Baonza, Antonio, and Matthew Freeman. 2002. “Control of Drosophila Eye Specification by Wingless Signalling.” Development (Cambridge, England) 129 (23): 5313–22. doi:10.1242/dev.00096.

Barolo, S, L A Carver, and J W Posakony. 2000. “GFP and Beta-Galactosidase Transformation Vectors for Promoter/enhancer Analysis in Drosophila.” BioTechniques 29 (4): 726, 728, 730, 732.

Baudry, Jean-Patrick, Cathy Maugis, Bertrand Michel, J.-P Baudry, and B Michel. 2012. “Slope Heuristics: Overview and Implementation.” Stat Comput 22: 455–70. doi:10.1007/s11222-011-9236-1.

Bauke, a.-C., S. Sasse, T. Matzat, and C. Klambt. 2015. “A Transcriptional Network Controlling Glial Development in the Drosophila Visual System.” Development 4 (May). highwire: 1–10. doi:10.1242/dev.119750.

Beckervordersandforth, Ruth M., Christof Rickert, Benjamin Altenhein, and Gerhard M. Technau. 2008. “Subtypes of Glial Cells in the Drosophila Embryonic Ventral Nerve Cord as Related to Lineage and Gene Expression.” Mechanisms of Development 125 (5–6): 542–57. doi:10.1016/j.mod.2007.12.004.

Bender, M., F.R. Turner, and T.C. Kaufman. 1987. “A Developmental Genetic Analysis of the Gene Regulator of Postbithorax in Drosophila Melanogaster.” Developmental Biology 119 (2): 418–32. doi:10.1016/0012-1606(87)90046-7.

Bogdan, Sven, and Christian Klämbt. 2001. “Epidermal Growth Factor Receptor Signaling.” Current Biology 11 (8): R292–95. doi:10.1016/S0960-9822(01)00167-1.

Brennan, C A, M. Ashburner, K. Moses, C. Antoniewski, B. Mugat, F. Delbac, J.-A. Lepesant, et al. 1998. “Ecdysone Pathway Is Required for Furrow Progression in the Developing Drosophila Eye.” Development (Cambridge, England) 125 (14): 2653–64. doi:10.1016/s0012-1606(74)80016-3.

Brennan, C A, T R Li, M Bender, F Hsiung, K Moses, J. Alcedo, M. Ayzenzon, et al. 2001. “Broad-Complex, but Not Ecdysone Receptor, Is Required for Progression of the Morphogenetic Furrow in the Drosophila Eye.” Development (Cambridge, England) 128 (1): 1–11. doi:10.1016/s0092-8674(00)80094-x.

Brown, N L, S W Paddock, C a Sattler, C Cronmiller, B J Thomas, and S B Carroll. 1996. “Daughterless Is Required for Drosophila Photoreceptor Cell Determination, Eye Morphogenesis, and Cell Cycle Progression.” Developmental Biology 179 (1): 65–78. doi:10.1006/dbio.1996.0241.

Campbell, G, and A Tomlinson. 1998. “The Roles of the Homeobox Genes Aristaless and Distal-Less in Patterning the Legs and Wings of Drosophila.” Development (Cambridge, England) 125 (22): 4483–93.

Carlson, Stanley D., Susan L. Hilgers, and Martin B. Garment. 1998. “Blood-Eye Barrier of the Developing Drosophila Melanogaster (Diptera: Drosophilidae).” International Journal of Insect Morphology and Embryology 27 (3): 241–47. doi:10.1016/S0020-7322(98)00016-6.

Carlson, Stanley D, Jyh-lyh Juang, Susan L Hilgers, and Martin B Garment. 2000. “Blood Barriers of the Insect.” Annual Review of Entomology 45 (1): 151–74. doi:10.1146/annurev.ento.45.1.151.

Casares, F, and R S Mann. 1998. “Control of Antennal versus Leg Development in Drosophila.” Nature 392 (6677): 723–26. doi:10.1038/33706.

Casares, Fernando, and Isabel Almudi. 2016. “Fast and Furious 800. The Retinal Determination Gene Network in Drosophila.” In Organogenetic Gene Networks, edited by James Castelli-Gair Hombría and Paola Bovolenta, 95–124. Cham: Springer International Publishing. doi:10.1007/978-3-319-42767-6.

Cavodeassi, F, R Diez Del Corral, S Campuzano, and M Domínguez. 1999. “Compartments and Organising Boundaries in the Drosophila Eye: The Role of the Homeodomain Iroquois Proteins.” Development (Cambridge, England) 126 (22): 4933–42.

Celniker, Susan E, Laura A L Dillon, Mark B Gerstein, Kristin C Gunsalus, Steven Henikoff, Gary H Karpen, Manolis Kellis, et al. 2009. “Unlocking the Secrets of the Genome.” Nature 459 (7249): 927–30. doi:10.1038/459927a.

Chen, R, G Halder, Z Zhang, and G Mardon. 1999. “Signaling by the TGF-Beta Homolog Decapentaplegic Functions Reiteratively within the Network of Genes Controlling Retinal Cell Fate Determination in Drosophila.” Development (Cambridge, England) 126 (5): 935–43.

Cheyette, Benjamin N R, Patricia J. Green, Kathy Martin, Hideki Garren, Volker Hartenstein, and S. Lawrence Zipursky. 1994. “The Drosophila Sine Oculis Locus Encodes a Homeodomain-Containing Protein Required for the Development of the Entire Visual System.” Neuron 12 (5): 977–96. doi:10.1016/0896-6273(94)90308-5.

Cho, K O, J Chern, S Izaddoost, and K W Choi. 2000. “Novel Signaling from the Peripodial Membrane Is Essential for Eye Disc Patterning in Drosophila.” Cell 103 (2): 331–42. doi:10.1016/S0092-8674(00)00124-0.

Choi, K. W., and S. Benzer. 1994. “Migration of Glia along Photoreceptor Axons in the Developing Drosophila Eye.” Neuron 12 (2): 423–31. doi:10.1016/0896-6273(94)90282-8.

Ciofani, Maria, Aviv Madar, Carolina Galan, MacLean Sellars, Kieran Mace, Florencia Pauli, Ashish Agarwal, et al. 2012. “A Validated Regulatory Network for Th17 Cell Specification.” Cell 151 (2). Elsevier Inc.: 289–303. doi:10.1016/j.cell.2012.09.016.

Cline, Melissa S, Michael Smoot, Ethan Cerami, Allan Kuchinsky, Nerius Landys, Chris Workman, Rowan Christmas, et al. 2007. “Integration of Biological Networks and Gene Expression Data Using Cytoscape.” Nature Protocols 2 (10): 2366–82. doi:10.1038/nprot.2007.324.

Cohen, Stephen M. 1993. “Imaginal Disc Development.” The Development of {IDrosophila Melanogaster} II. Cold Spring Harbor Laboratory Press: 747–841.

Consortium, The Gene Ontology. 2015. “Gene Ontology Consortium: Going Forward.” Nucleic Acids Research 43 (D1). Oxford University Press: D1049–56. doi:10.1093/nar/gku1179.

Cummins, Mark, Jose I. Pueyo, Steve A. Greig, and Juan Pablo Couso. 2003. “Comparative Analysis of Leg and Antenna Development in Wild-Type and Homeotic Drosophila Melanogaster.” Development Genes and Evolution 213 (7): 319–27. doi:10.1007/s00427-003-0326-8.

Dai, P, H Akimaru, Y Tanaka, D X Hou, T Yasukawa, C Kanei-Ishii, T Takahashi, and S Ishii. 1996. “CBP as a Transcriptional Coactivator of c-Myb.” Genes & Development 10 (5): 528–40.

Daines, Bryce, H. Wang, L. Wang, Y. Li, Yi Han, David Emmert, William Gelbart, et al. 2011. “The Drosophila Melanogaster Transcriptome by Paired-End RNA Sequencing.” Genome Research 21 (2): 315–24. doi:10.1101/gr.107854.110.

Deplancke, Bart, Arnab Mukhopadhyay, Wanyuan Ao, Ahmed M. Elewa, Christian A. Grove, Natalia J. Martinez, Reynaldo Sequerra, et al. 2006. “A Gene-Centered. C. Elegans Protein-DNA Interaction Network.” Cell 125 (6): 1193–1205. doi:10.1016/j.cell.2006.04.038.

DeSalvo, Michael K., Samantha J. Hindle, Zeid M. Rusan, Souvinh Orng, Mark Eddison, Kyle Halliwill, and Roland J. Bainton. 2014. “The Drosophila Surface Glia Transcriptome: Evolutionary Conserved Blood-Brain Barrier Processes.” Frontiers in Neuroscience 8 (November): 1–22. doi:10.3389/fnins.2014.00346.

Dey, Bijan Kumar, Xiao-Li Zhao, Emmanuel Popo-Ola, and Ana Regina Campos. 2009. “Mutual Regulation of the Drosophila Disconnected (Disco) and Distal-Less (Dll) Genes Contributes to Proximal-Distal Patterning of Antenna and Leg.” Cell and Tissue Research 338 (2): 227–40. doi:10.1007/s00441-009-0865-z.

Domínguez, M. 1999. “Dual Role for Hedgehog in the Regulation of the Proneural Gene Atonal during Ommatidia Development.” Development (Cambridge, England) 126 (11): 2345–53.

Domínguez, María, and Fernando Casares. 2005. “Organ Specification-Growth Control Connection: New in-Sights from the Drosophila Eye-Antennal Disc.” Developmental Dynamics 232 (3): 673–84. doi:10.1002/dvdy.20311.

Domínguez, María, and Ernst Hafen. 1997. “Hedgehog Directly Controls Initiation and Propagation of Retinal Differentiation in the Drosophila Eye.” Genes and Development 11 (23): 3254–64. doi:10.1101/gad.11.23.3254.

Dong, P D, J Chu, and G Panganiban. 2000. “Coexpression of the Homeobox Genes Distal-Less and Homothorax Determines Drosophila Antennal Identity.” Development (Cambridge, England) 127 (2): 209–16.

Dong, P D Si, Jennifer Scholz Dicks, and Grace Panganiban. 2002. “Distal-Less and Homothorax Regulate Multiple Targets to Pattern the Drosophila Antenna.” Development (Cambridge, England) 129 (8): 1967–74.

Edwards, J S, L S Swales, and M Bate. 1993. “The Differentiation between Neuroglia and Connective Tissue Sheath in Insect Ganglia Revisited: The Neural Lamella and Perineurial Sheath Cells Are Absent in a Mesodermless Mutant of Drosophila.” The Journal of Comparative Neurology 333 (2): 301–8. doi:10.1002/cne.903330214.

Edwards, Tara N., and Ian a. Meinertzhagen. 2010. “The Functional Organisation of Glia in the Adult Brain of Drosophila and Other Insects.” Progress in Neurobiology 90 (4). Elsevier Ltd: 471–97. doi:10.1016/j.pneurobio.2010.01.001.

Edwards, Tara N, Andrea C Nuschke, Aljoscha Nern, and Ian A Meinertzhagen. 2012. “Organization and Metamorphosis of Glia in the Drosophila Visual System.” The Journal of Comparative Neurology 520 (10): 2067–85. doi:10.1002/cne.23071.

Elliott, Jimmy, Christine Jolicoeur, Vasanth Ramamurthy, and Michel Cayouette. 2008. “Ikaros Confers Early Temporal Competence to Mouse Retinal Progenitor Cells.” Neuron 60 (1): 26–39. doi:10.1016/j.neuron.2008.08.008.

Frankfort, Benjamin J, and Graeme Mardon. 2002. “R8 Development in the Drosophila Eye: A Paradigm for Neural Selection and Differentiation.” Development (Cambridge, England) 129 (6): 1295–1306.

Franzdóttir, Sigrídur Rut, Daniel Engelen, Yeliz Yuva-Aydemir, Imke Schmidt, Annukka Aho, and Christian Klämbt. 2009. “Switch in FGF Signalling Initiates Glial Differentiation in the Drosophila Eye.” Nature 460 (7256). Macmillan Publishers Limited. All rights reserved: 758–61. doi:10.1038/nature08167.

Gao, Haixia, Xingjuan Chen, Xiaona Du, Bingcai Guan, Yani Liu, and Hailin Zhang. 2011. “EGF Enhances the Migration of Cancer Cells by up-Regulation of TRPM7.” Cell Calcium 50 (6). Elsevier Ltd: 559–68. doi:10.1016/j.ceca.2011.09.003.

Garcia-Bellido, a, and J R Merriam. 1969. “Cell Lineage of the Imaginal Discs in Drosophila Gynandromorphs.” The Journal of Experimental Zoology 170 (1): 61–75. doi:10.1002/jez.1401700106.

Grosskortenhaus, Ruth, Bret J Pearson, Amanda Marusich, and Chris Q Doe. 2005. “Regulation of Temporal Identity Transitions in Drosophila Neuroblasts.” Developmental Cell 8 (2): 193–202. doi:10.1016/j.devcel.2004.11.019.

Gunthorpe, D, K E Beatty, and M V Taylor. 1999. “Different Levels, but Not Different Isoforms, of the Drosophila Transcription Factor DMEF2 Affect Distinct Aspects of Muscle Differentiation.” Developmental Biology 215 (1): 130–45. doi:10.1006/dbio.1999.9449.

Heberlein, U, C M Singh, a Y Luk, and T J Donohoe. 1995. “Growth and Differentiation in the Drosophila Eye Coordinated by Hedgehog.” Nature 373 (6516): 709–11. doi:10.1038/373709a0.

Heberlein, U, and J E Treisman. 2000. “Early Retinal Development in Drosophila.” Results and Problems in Cell Differentiation 31: 37–50.

Heinz, Sven, Christopher Benner, Nathanael Spann, Eric Bertolino, Yin C. Lin, Peter Laslo, Jason X. Cheng, Cornelis Murre, Harinder Singh, and Christopher K. Glass. 2010. “Simple Combinations of Lineage-Determining Transcription Factors Prime Cis-Regulatory Elements Required for Macrophage and B Cell Identities.” Molecular Cell 38 (4): 576–89. doi:10.1016/j.molcel.2010.05.004.

Herboso, Leire, Marisa M Oliveira, Ana Talamillo, Coralia Pérez, Monika González, David Martín, James D Sutherland, Alexander W Shingleton, Christen K Mirth, and Rosa Barrio. 2015. “Ecdysone Promotes Growth of Imaginal Discs through the Regulation of Thor in D. Melanogaster.” Scientific Reports 5 (July). Nature Publishing Group: 12383. doi:10.1038/srep12383.

Herrmann, Carl, Bram Van de Sande, Delphine Potier, and Stein Aerts. 2012. “I-cisTarget: An Integrative Genomics Method for the Prediction of Regulatory Features and Cis-Regulatory Modules.” Nucleic Acids Research 40 (15): e114. doi:10.1093/nar/gks543.

Homem, C. C. F., and J. A. Knoblich. 2012. “Drosophila Neuroblasts: A Model for Stem Cell Biology.” Development 139 (23): 4297–4310. doi:10.1242/dev.080515.

Hummel, Thomas, Suzanne Attix, Dorian Gunning, and S. Lawrence Lawrence Zipursky. 2002. “Temporal Control of Glial Cell Migration in the Drosophila Eye Requires Gilgamesh, Hedgehog, and Eye Specification Genes.” Neuron 33 (2): 193–203. doi:10.1016/S0896-6273(01)00581-5.

Iadecola, Costantino, and Maiken Nedergaard. 2007. “Glial Regulation of the Cerebral Microvasculature.” Nature Neuroscience 10 (11): 1369–76. doi:10.1038/nn2003.

Ideker, Trey, V Thorsson, J A Ranish, R Christmas, J Buhler, J K Eng, R Bumgarner, D R Goodlett, R Aebersold, and L Hood. 2001. “Integrated Genomic and Proteomic Analyses of a Systematically Perturbed Metabolic Network.” Science (New York, N.Y.) 292 (5518): 929–34. doi:10.1126/science.292.5518.929.

Isshiki, Takako, Bret Pearson, Scott Holbrook, and Chris Q Doe. 2001. “Drosophila Neuroblasts Sequentially Express Transcription Factors Which Specify the Temporal Identity of Their Neuronal Progeny.” Cell 106 (4): 511–21. doi:10.1016/S0092-8674(01)00465-2.

Ito, Kei, Joachim Urban, and Gerhard Martin Technau. 1995. “Distribution, Classification, and Development ofDrosophila Glial Cells in the Late Embryonic and Early Larval Ventral Nerve Cord.” Roux’s Archives of Developmental Biology 204 (5): 284–307. doi:10.1007/BF02179499.

Jarman, a P, E H Grell, L Ackerman, L Y Jan, and Y N Jan. 1994. “Atonal Is the Proneural Gene for Drosophila Photoreceptors.” Nature 369 (6479): 398–400. doi:10.1038/369398a0.

Jarman, a P, Y Sun, L Y Jan, and Y N Jan. 1995. “Role of the Proneural Gene, Atonal, in Formation of Drosophila Chordotonal Organs and Photoreceptors.” Development (Cambridge, England) 121 (7): 2019–30.

Jarvis, Lesley a. 2006. “Sprouty Proteins Are in Vivo Targets of Corkscrew/SHP-2 Tyrosine Phosphatases.” Development 133 (6): 1133–42. doi:10.1242/dev.02255.

Jiménez, Fernando, and JoséA. Campos-Ortega. 1990. “Defective Neuroblast Commitment in Mutants of the Achaete-Scute Complex and Adjacent Genes of D. Melanogaster.” Neuron 5 (1): 81–89. doi:10.1016/0896-6273(90)90036-F.

Jockusch, Elizabeth L, and Frank W Smith. 2015. “Hexapoda: Comparative Aspects of Later Embryogenesis and Metamorphosis.” In Evolutionary Developmental Biology of Invertebrates 5: Ecdysozoa III: Hexapoda, edited by Andreas Wanninger, 111–208. Vienna: Springer Vienna. doi:10.1007/978-3-7091-1868-9_3.

Junion, Guillaume, Mikhail Spivakov, Charles Girardot, Martina Braun, E. Hilary Gustafson, Ewan Birney, and Eileen E. M. E. E. E M. Furlong. 2012. “A Transcription Factor Collective Defines Cardiac Cell Fate and Reflects Lineage History.” Cell 148 (3): 473–86. doi:10.1016/j.cell.2012.01.030.

Jusiak, Barbara, Umesh C Karandikar, Su-Jin Kwak, Feng Wang, Hui Wang, Rui Chen, and Graeme Mardon. 2014. “Regulation of Drosophila Eye Development by the Transcription Factor Sine Oculis.” Edited by Justin Kumar. PloS One 9 (2): e89695. doi:10.1371/journal.pone.0089695.

Juven-Gershon, Tamar, Jer-Yuan Hsu, and James T Kadonaga. 2008. “Caudal, a Key Developmental Regulator, Is a DPE-Specific Transcriptional Factor.” Genes & Development 22 (20): 2823–30. doi:10.1101/gad.1698108.

Kambadur, Ravi, Keita Koizumi, Chad Stivers, James Nagle, Stephen J. Poole, and Ward F. Odenwald. 1998. “Regulation of POU Genes by Castor and Hunchback Establishes Layered Compartments in the Drosophila CNS.” Genes & Development 12 (2): 246–60. doi:10.1101/gad.12.2.246.

Kango-Singh, Madhuri, Amit Singh, and Y. Henry Sun. 2003. “Eyeless Collaborates with Hedgehog and Decapentaplegic Signaling in Drosophila Eye Induction.” Developmental Biology 256 (1): 48–60. doi:10.1016/S0012-1606(02)00123-9.

Kemmeren, Patrick, Katrin Sameith, Loes A.L. van de Pasch, Joris J. Benschop, Tineke L. Lenstra, Thanasis Margaritis, Eoghan O‘Duibhir, et al. 2014. “Large-Scale Genetic Perturbations Reveal Regulatory Networks and an Abundance of Gene-Specific Repressors.” Cell 157 (3). Elsevier: 740–52. doi:10.1016/j.cell.2014.02.054.

Kent, W James, Charles W Sugnet, Terrence S Furey, Krishna M Roskin, Tom H Pringle, Alan M Zahler, and David Haussler. 2002. “The Human Genome Browser at UCSC.” Genome Research 12 (6): 996–1006. doi:10.1101/gr.229102.

Kenyon, Kristy L., Swati S. Ranade, Jennifer Curtiss, Marek Mlodzik, and Francesca Pignoni. 2003. “Coordinating Proliferation and Tissue Specification to Promote Regional Identity in the Drosophila Head.” Developmental Cell 5 (3): 403–14. doi:10.1016/S1534-5807(03)00243-0.

Kim, Hong Joo, and Dafna Bar-Sagi. 2004. “Modulation of Signalling by Sprouty: A Developing Story.” Nature Reviews Molecular Cell Biology 5 (6): 441–50. doi:10.1038/nrm1400.

Klein, Thomas. 2008. “Immunolabeling of Imaginal Discs.” Methods in Molecular Biology (Clifton, N.J.) 420. Humana Press Inc.: 253–63. doi:10.1007/978-1-59745-583-1_15.

Koestler, Stefan A, Begum Alaybeyoglu, Christian X Weichenberger, and Arzu Celik. 2015. “FlyOde - a Platform for Community Curation and Interactive Visualization of Dynamic Gene Regulatory Networks in Drosophila Eye Development [Version 1; Referees: Awaiting Peer Review].” F1000Research 4 (1): 1–6. doi:10.12688/f1000research.7556.1.

Kosman, David, Stephen Small, and J. Reinitz. 1998. “Rapid Preparation of a Panel of Polyclonal Antibodies to Drosophila Segmentation Proteins.” Development Genes and Evolution 208 (5): 290–94. doi:10.1007/s004270050184.

Kramer, S, M Okabe, N Hacohen, M A Krasnow, and Y Hiromi. 1999. “Sprouty: A Common Antagonist of FGF and EGF Signaling Pathways in Drosophila.” Development (Cambridge, England) 126 (11): 2515–25.

Kumar, Arun, Tripti Gupta, Sara Berzsenyi, Angela Giangrande, B. Aigouy, V. Van de Bor, M. Boeglin, et al. 2015. “N-Cadherin Negatively Regulates Collective Drosophila Glial Migration through Actin Cytoskeleton Remodeling.” Journal of Cell Science 128 (5). The Company of Biologists Ltd: 900–912. doi:10.1242/jcs.157974.

Kumar, J P, and K Moses. 2001. “EGF Receptor and Notch Signaling Act Upstream of Eyeless/Pax6 to Control Eye Specification.” Cell 104 (5): 687–97. doi:10.1016/S0092-8674(02)08063-7.

Kumar, Justin P, Tazeen Jamal, Alex Doetsch, F Rudolf Turner, and Joseph B Duffy. 2004. “CREB Binding Protein Functions during Successive Stages of Eye Development in Drosophila.” Genetics 168 (2): 877–93. doi:10.1534/genetics.104.029850.

Kurata, S, M J Go, S Artavanis-Tsakonas, and W J Gehring. 2000. “Notch Signaling and the Determination of Appendage Identity.” Proceedings of the National Academy of Sciences of the United States of America 97 (5): 2117–22. doi:10.1073/pnas.040556497.

Kwok, R P, J R Lundblad, J C Chrivia, J P Richards, H P Bächinger, R G Brennan, S G Roberts, M R Green, and R H Goodman. 1994. “Nuclear Protein CBP Is a Coactivator for the Transcription Factor CREB.” Nature 370 (6486): 223–26. doi:10.1038/370223a0.

Langmead, Ben, and Steven L Salzberg. 2012. “Fast Gapped-Read Alignment with Bowtie 2.” Nature Methods 9 (4): 357–59. doi:10.1038/nmeth.1923.

Lebreton, Gaëlle, Christian Faucher, David L Cribbs, and Corinne Benassayag. 2008. “Timing of Wingless Signalling Distinguishes Maxillary and Antennal Identities in Drosophila Melanogaster.” Development (Cambridge, England) 135 (13): 2301–9. doi:10.1242/dev.017053.

Lee, Tong Ihn, Nicola J Rinaldi, François Robert, Duncan T Odom, Ziv Bar-Joseph, Georg K Gerber, Nancy M Hannett, et al. 2002. “Transcriptional Regulatory Networks in Saccharomyces Cerevisiae.” Science (New York, N.Y.) 298 (5594): 799–804. doi:10.1126/science.1075090.

Lehmann, R, and C Nüsslein-Volhard. 1987. “Hunchback, a Gene Required for Segmentation of an Anterior and Posterior Region of the Drosophila Embryo.” Developmental Biology 119 (2): 402–17. doi:10.1016/0012-1606(87)90045-5.

Li, Heng, Bob Handsaker, Alec Wysoker, Tim Fennell, Jue Ruan, Nils Homer, Gabor Marth, Goncalo Abecasis, and Richard Durbin. 2009. “The Sequence Alignment/Map Format and SAMtools.” Bioinformatics 25 (16): 2078–79. doi:10.1093/bioinformatics/btp352.

Li, T, and M Bender. 2000. “A Conditional Rescue System Reveals Essential Functions for the Ecdysone Receptor (EcR) Gene during Molting and Metamorphosis in Drosophila.” Development (Cambridge, England) 127 (13): 2897–2905.

Li, Xiao-yong, Stewart MacArthur, Richard Bourgon, David Nix, Daniel A Pollard, Venky N Iyer, Aaron Hechmer, et al. 2008. “Transcription Factors Bind Thousands of Active and Inactive Regions in the Drosophila Blastoderm.” Edited by Jim Kadonaga. PLoS Biology 6 (2). Public Library of Science: e27. doi:10.1371/journal.pbio.0060027.

Lilly, B, S Galewsky, A B Firulli, R A Schulz, and E N Olson. 1994. “D-MEF2: A MADS Box Transcription Factor Expressed in Differentiating Mesoderm and Muscle Cell Lineages during Drosophila Embryogenesis.” Proceedings of the National Academy of Sciences of the United States of America 91 (12): 5662–66.

Love, M. I., Simon Anders, and Wolfgang Huber. 2014. “Differential Analysis of Count Data - the DESeq2 Package,” 1–41. doi:10.1101/002832.

MacArthur, Stewart, Xiao-Yong Li, Jingyi Li, James B Brown, Hou Cheng Chu, Lucy Zeng, Brandi P Grondona, et al. 2009. “Developmental Roles of 21 Drosophila Transcription Factors Are Determined by Quantitative Differences in Binding to an Overlapping Set of Thousands of Genomic Regions.” Genome Biology 10 (7): R80. doi:10.1186/gb-2009-10-7-r80.

MacManes, Matthew D. 2014. “On the Optimal Trimming of High-Throughput mRNA Sequence Data.” Frontiers in Genetics 5 (JAN): 1–7. doi:10.3389/fgene.2014.00013.

Maere, Steven, Karel Heymans, and Martin Kuiper. 2005. “BiNGO: A Cytoscape Plugin to Assess Overrepresentation of Gene Ontology Categories in Biological Networks.” Bioinformatics (Oxford, England) 21 (16): 3448–49. doi:10.1093/bioinformatics/bti551.

Malartre, Marianne. 2016. “Regulatory Mechanisms of EGFR Signalling during Drosophila Eye Development.” Cellular and Molecular Life Sciences, March. Springer International Publishing. doi:10.1007/s00018-016-2153-x.

Mao, Yanlan, and Matthew Freeman. 2009. “Fasciclin 2, the Drosophila Orthologue of Neural Cell-Adhesion Molecule, Inhibits EGF Receptor Signalling.” Development (Cambridge, England) 136 (3): 473–81. doi:10.1242/dev.026054.

Mardon, G, N M Solomon, and G M Rubin. 1994. “Dachshund Encodes a Nuclear Protein Required for Normal Eye and Leg Development in Drosophila.” Development (Cambridge, England) 120 (12): 3473–86.

Maurel-Zaffran, C, and J E Treisman. 2000. “Pannier Acts Upstream of Wingless to Direct Dorsal Eye Disc Development in Drosophila.” Development (Cambridge, England) 127 (5): 1007–16.

McManus, K J, and M J Hendzel. 2001. “CBP, a Transcriptional Coactivator and Acetyltransferase.” Biochemistry and Cell Biology = Biochimie et Biologie Cellulaire 79 (3): 253–66.

Merrill, V.K.L., F.R. Turner, and T.C. Kaufman. 1987. “A Genetic and Developmental Analysis of Mutations in the Deformed Locus in Drosophila Melanogaster.” Developmental Biology 122 (2): 379–95. doi:10.1016/0012-1606(87)90303-4.

Merrill, V K, R J Diederich, F R Turner, and T C Kaufman. 1989. “A Genetic and Developmental Analysis of Mutations in Labial, a Gene Necessary for Proper Head Formation in Drosophila Melanogaster.” Developmental Biology 135 (2): 376–91. doi:10.1016/0012-1606(89)90187-5.

Mishra, Abhishek Kumar, Bastiaan O.R. Bargmann, Maria Tsachaki, Cornelia Fritsch, and Simon G. Sprecher. 2016. “Functional Genomics Identifies Regulators of the Phototransduction Machinery in the Drosophila Larval Eye and Adult Ocelli.” Developmental Biology 410 (2). Elsevier: 164–77. doi:10.1016/j.ydbio.2015.12.026.

Moberg, Kenneth H, Daphne W Bell, Doke C. R. Wahrer, Daniel A Haber, and Iswar K Hariharan. 2001. “Archipelago Regulates Cyclin E Levels in Drosophila and Is Mutated in Human Cancer Cell Lines.” Nature 413 (6853): 311–16. doi:10.1038/35095068.

Morata, G. 2001. “How Drosophila Appendages Develop.” Nature Reviews. Molecular Cell Biology 2 (2): 89–97. doi:10.1038/35052047.

Naval-Sańchez, Marina, Delphine Potier, Lotte Haagen, Máximo Sánchez, Sebastian Munck, Bram Van De Sande, Fernando Casares, Valerie Christiaens, and Stein Aerts. 2013. “Comparative Motif Discovery Combined with Comparative Transcriptomics Yields Accurate Targetome and Enhancer Predictions.” Genome Research 23 (1): 74–88. doi:10.1101/gr.140426.112.

Nègre, Nicolas, Christopher D Brown, Lijia Ma, Christopher Aaron Bristow, Steven W Miller, Ulrich Wagner, Pouya Kheradpour, et al. 2011. “A Cis-Regulatory Map of the Drosophila Genome.” Nature 471 (7339): 527–31. doi:10.1038/nature09990.

Nguyen, H T, R Bodmer, S M Abmayr, J C McDermott, and N A Spoerel. 1994. “D-mef2: A Drosophila Mesoderm-Specific MADS Box-Containing Gene with a Biphasic Expression Profile during Embryogenesis.” Proceedings of the National Academy of Sciences of the United States of America 91 (16): 7520–24.

Nüsslein-Volhard, Christiane, and Eric Wieschaus. 1980. “Mutations Affecting Segment Number and Polarity in Drosophila.” Nature 287 (5785). Nature Publishing Group: 795–801. doi:10.1038/287795a0.

O’Neill, Elizabeth M., Ilaria Rebay, Robert Tjian, and Gerald M. Rubin. 1994. “The Activities of Two Ets-Related Transcription Factors Required for Drosophila Eye Development Are Modulated by the Ras/MAPK Pathway.” Cell 78 (1): 137–47. doi:10.1016/0092-8674(94)90580-0.

Oros, Sarah M., Meghana Tare, Madhuri Kango-Singh, and Amit Singh. 2010. “Dorsal Eye Selector Pannier (Pnr) Suppresses the Eye Fate to Define Dorsal Margin of the Drosophila Eye.” Developmental Biology 346 (2): 258–71. doi:10.1016/j.ydbio.2010.07.030.

Petrif, Fred, Rachel Giles, H., Hans Dauwerse, G., Jasper Saris, J., et al. 1995. “Rubinstein-Taybi Syndrome Caused by Mutations in the Transcriptional Co-Activator CBP.” Nature 376 (6538): 348–51. doi:10.1038/376348a0.

Pfeiffer, Barret D, Arnim Jenett, Ann S Hammonds, Teri-T B Ngo, Sima Misra, Christine Murphy, Audra Scully, et al. 2008. “Tools for Neuroanatomy and Neurogenetics in Drosophila.” Proceedings of the National Academy of Sciences of the United States of America 105 (28): 9715–20. doi:10.1073/pnas.0803697105.

Pichaud, Franck, and Fernando Casares. 2000. “Homothorax and Iroquois-C Genes Are Required for the Establishment of Territories within the Developing Eye Disc.” Mechanisms of Development 96 (1): 15–25. doi:10.1016/S0925-4773(00)00372-5.

Pinnell, Jamie. 2006. “The Divergent Roles of the Segmentation Gene Hunchback.” Integrative and Comparative Biology 46 (4): 519–32. doi:10.1093/icb/icj054.

Pinsonneault, Robert L, Nasima Mayer, Fahima Mayer, Nebiyu Tegegn, and Roland J Bainton. 2011. “Novel Models for Studying the Blood-Brain and Blood-Eye Barriers in Drosophila.” Methods in Molecular Biology (Clifton, N.J.) 686: 357–69. doi:10.1007/978-1-60761-938-3_17.

Posnien, Nico, Corinna Hopfen, Maarten Hilbrant, Margarita Ramos-Womack, Sophie Murat, Anna Schönauer, Samantha L. Herbert, et al. 2012. “Evolution of Eye Morphology and Rhodopsin Expression in the Drosophila Melanogaster Species Subgroup.” PLoS ONE 7 (5). doi:10.1371/journal.pone.0037346.

Potier, Delphine, Kristofer Davie, Gert Hulselmans, Marina Naval Sanchez, Lotte Haagen, Vân Anh Huynh-Thu, Duygu Koldere, et al. 2014. “Mapping Gene Regulatory Networks in Drosophila Eye Development by Large-Scale Transcriptome Perturbations and Motif Inference.” Cell Reports 9 (6): 2290–2303. doi:10.1016/j.celrep.2014.11.038.

Quiring, R, U Walldorf, U Kloter, and W J Gehring. 1994. “Homology of the Eyeless Gene of Drosophila to the Small Eye Gene in Mice and Aniridia in Humans.” Science (New York, N.Y.) 265 (5173): 785–89. doi:10.1126/science.7914031.

Rangarajan, Radha, Hélène Courvoisier, and Ulrike Gaul. 2001. “Dpp and Hedgehog Mediate Neuron-Glia Interactions in Drosophila Eye Development by Promoting the Proliferation and Motility of Subretinal Glia.” Mechanisms of Development 108 (1–2): 93–103. doi:10.1016/S0925-4773(01)00501-9.

Rangarajan, R, Q Gong, and U Gaul. 1999. “Migration and Function of Glia in the Developing Drosophila Eye.” Development (Cambridge, England) 126 (15): 3285–92.

Rau, Andrea, Mélina Gallopin, Gilles Celeux, and Florence Jaffrézic. 2013. “Data-Based Filtering for Replicated High-Throughput Transcriptome Sequencing Experiments.” Bioinformatics (Oxford, England) 29 (17): 2146–52. doi:10.1093/bioinformatics/btt350.

Rau, Andrea, Cathy Maugis-Rabusseau, Marie-Laure Martin-Magniette, and Gilles Celeux. 2015. “Co-Expression Analysis of High-Throughput Transcriptome Sequencing Data with Poisson Mixture Models.” Bioinformatics (Oxford, England) 31 (January): 1420–27. doi:10.1093/bioinformatics/btu845.

Reichert, Heinrich. 2011. “Drosophila Neural Stem Cells: Cell Cycle Control of Self-Renewal, Differentiation, and Termination in Brain Development.” In Results and Problems in Cell Differentiation, edited by Jacek Z. Kubiak, 53:229–46. Results and Problems in Cell Differentiation. Berlin, Heidelberg: Springer Berlin Heidelberg. doi:10.1007/978-3-642-19065-0.

Richardson, Brian E, and Ruth Lehmann. 2010. “Mechanisms Guiding Primordial Germ Cell Migration: Strategies from Different Organisms.” Nature Reviews Molecular Cell Biology 11 (1). Nature Publishing Group: 37–49. doi:10.1038/nrm2815.

Rochlin, Kate, Shannon Yu, Sudipto Roy, and Mary K. Baylies. 2010. “Myoblast Fusion: When It Takes More to Make One.” Developmental Biology 341 (1). Elsevier Inc.: 66–83. doi:10.1016/j.ydbio.2009.10.024.

Roy, Sushmita, Jason Ernst, Peter V Kharchenko, Pouya Kheradpour, Nicolas Negre, Matthew L Eaton, Jane M Landolin, et al. 2010. “Identification of Functional Elements and Regulatory Circuits by Drosophila modENCODE.” Science (New York, N.Y.) 330 (6012): 1787–97. doi:10.1126/science.1198374.

Sato, Atsushi, and Andrew Tomlinson. 2007. “Dorsal-Ventral Midline Signaling in the Developing Drosophila Eye.” Development (Cambridge, England) 134 (4): 659–67. doi:10.1242/dev.02786.

Schröder, Reinhard. 2003. “The Genes Orthodenticle and Hunchback Substitute for Bicoid in the Beetle Tribolium.” Nature 422 (April): 621–25. doi:10.1038/nature01518.1.

Schwabe, Tina, Roland J. Bainton, Richard D. Fetter, Ulrike Heberlein, and Ulrike Gaul. 2005. “GPCR Signaling Is Required for Blood-Brain Barrier Formation in Drosophila.” Cell 123 (1): 133–44. doi:10.1016/j.cell.2005.08.037.

Serikaku, M. a., and J. E. O’Tousa. 1994. “Sine Oculis Is a Homeobox Gene Required for Drosophila Visual System Development.” Genetics 138 (4): 1137–50.

Sharma, H S, and P K Dey. 1986. “Influence of Long-Term Immobilization Stress on Regional Blood-Brain Barrier Permeability, Cerebral Blood Flow and 5-HT Level in Conscious Normotensive Young Rats.” Journal of the Neurological Sciences 72 (1): 61–76.

Sharma, Sreenath V, Daphne W Bell, Jeffrey Settleman, and Daniel A Haber. 2007. “Epidermal Growth Factor Receptor Mutations in Lung Cancer.” Nature Reviews Cancer 7 (3): 169–81. doi:10.1038/nrc2088.

Shcherbata, Halyna R. 2004. “The Mitotic-to-Endocycle Switch in Drosophila Follicle Cells Is Executed by Notch-Dependent Regulation of G1/S, G2/M and M/G1 Cell-Cycle Transitions.” Development 131 (13): 3169–81. doi:10.1242/dev.01172.

Shen, W, and G Mardon. 1997. “Ectopic Eye Development in Drosophila Induced by Directed Dachshund Expression.” Development (Cambridge, England) 124 (1): 45–52.

Shilo, B. 2003. “Signaling by the Drosophila Epidermal Growth Factor Receptor Pathway during Development.” Experimental Cell Research 284 (1): 140–49. doi:10.1016/S0014-4827(02)00094-0.

Shilo, B.-Z. 2005. “Regulating the Dynamics of EGF Receptor Signaling in Space and Time.” Development 132 (18): 4017–27. doi:10.1242/dev.02006.

Shir-Shapira, Hila, Julia Sharabany, Matan Filderman, Diana Ideses, Avital Ovadia-Shochat, Mattias Mannervik, and Tamar Juven-Gershon. 2015. “Structure-Function Analysis of the Drosophila Melanogaster Caudal Transcription Factor Provides Insights into Core Promoter-Preferential Activation.” The Journal of Biological Chemistry 290 (28): 17293–305. doi:10.1074/jbc.M114.632109.

Sieglitz, Florian, Till Matzat, Yeliz Yuva-Aydemir, Helen Neuert, Benjamin Altenhein, and C. Klambt. 2013. “Antagonistic Feedback Loops Involving Rau and Sprouty in the Drosophila Eye Control Neuronal and Glial Differentiation.” Science Signaling 6 (300): ra96-ra96. doi:10.1126/scisignal.2004651.

Silies, Marion, Yeliz Yuva, Daniel Engelen, Annukka Aho, Tobias Stork, and Christian Klämbt. 2007. “Glial Cell Migration in the Eye Disc.” The Journal of Neuroscience: The Official Journal of the Society for Neuroscience 27 (48): 13130–39. doi:10.1523/JNEUROSCI.3583-07.2007.

Singh, Amit, Jeeder Chan, Joshua J Chern, and Kwang-Wook Choi. 2005. “Genetic Interaction of Lobe with Its Modifiers in Dorsoventral Patterning and Growth of the Drosophila Eye.” Genetics 171 (1): 169–83. doi:10.1534/genetics.105.044180.

Singh, Amit, and Kwang-Wook Choi. 2003. “Initial State of the Drosophila Eye before Dorsoventral Specification Is Equivalent to Ventral.” Development (Cambridge, England) 130 (25): 6351–60. doi:10.1242/dev.00864.

Sivachenko, Anna, Yue Li, Katharine C Abruzzi, and Michael Rosbash. 2013. “The Transcription Factor Mef2 Links the Drosophila Core Clock to Fas2, Neuronal Morphology, and Circadian Behavior.” Neuron 79 (2): 281–92. doi:10.1016/j.neuron.2013.05.015.

Skeath, James B., and Stefan Thor. 2003. “Genetic Control of Drosophila Nerve Cord Development.” Current Opinion in Neurobiology 13 (1): 8–15. doi:10.1016/S0959-4388(03)00007-2.

Skultétyová, I, D Tokarev, and D Jezová. 1998. “Stress-Induced Increase in Blood-Brain Barrier Permeability in Control and Monosodium Glutamate-Treated Rats.” Brain Research Bulletin 45 (2): 175–78.

Snodgrass, Robert E. 1935. “Principles of Insect Morphology” 136 (3447). Ithaca, United States: {McGraw-Hill} Book Co: 812–13. http://www.nature.com/doifinder/10.1038/136812a0.

Tautz, Diethard, Ruth Lehmann, Harald Schnurch, Reinhard Schuh, Eveline Seifert, Andrea Kienlin, Keith Jones, and Herbert Jackle. 1987. “Finger Protein of Novel Structure Encoded by Hunchback, a Second Member of the Gap Class of Drosophila Segmentation Genes.” Nature 327 (6121): 383–89.

Tavazoie, Saeed, Jason D Hughes, Michael J Campbell, Raymond J Cho, and George M Church. 1999. “Systematic Determination of Genetic Network Architecture.” Nature Genetics 22 (3): 281–85. doi:10.1038/10343.

Treisman, J E, and U Heberlein. 1998. “Eye Development in Drosophila: Formation of the Eye Field and Control of Differentiation.” Current Topics in Developmental Biology 39: 119–58.

Treisman, Jessica E. 2013. “Retinal Differentiation in Drosophila.” Wiley Interdisciplinary Reviews: Developmental Biology 2 (4): 545–57. doi:10.1002/wdev.100.

Unhavaithaya, Yingdee, and T. L. Orr-Weaver. 2012. “Polyploidization of Glia in Neural Development Links Tissue Growth to Blood-Brain Barrier Integrity.” Genes & Development 26 (1): 31–36. doi:10.1101/gad.177436.111.

van Genderen, M M, G F Kinds, F C Riemslag, and R C Hennekam. 2000. “Ocular Features in Rubinstein-Taybi Syndrome: Investigation of 24 Patients and Review of the Literature.” The British Journal of Ophthalmology 84 (10): 1177–84. doi:10.1136/bjo.84.10.1177.

von Hilchen, Christian M, Alvaro E Bustos, Angela Giangrande, Gerhard M Technau, and Benjamin Altenhein. 2013. “Predetermined Embryonic Glial Cells Form the Distinct Glial Sheaths of the Drosophila Peripheral Nervous System.” Development (Cambridge, England) 140 (17): 3657–68. doi:10.1242/dev.093245.

Weber, Ursula, Csilla Pataki, Jozsef Mihaly, and Marek Mlodzik. 2008. “Combinatorial Signaling by the Frizzled/PCP and Egfr Pathways during Planar Cell Polarity Establishment in the Drosophila Eye.” Developmental Biology 316 (1): 110–23. doi:10.1016/j.ydbio.2008.01.016.

Williams, Claire R, Alyssa Baccarella, Jay Z Parrish, and Charles C Kim. 2016. “Trimming of Sequence Reads Alters RNA-Seq Gene Expression Estimates.” BMC Bioinformatics 17 (1). BMC Bioinformatics: 103. doi:10.1186/s12859-016-0956-2.

Wolff, T, and D F Ready. 1991. “The Beginning of Pattern Formation in the Drosophila Compound Eye: The Morphogenetic Furrow and the Second Mitotic Wave.” Development (Cambridge, England) 113 (3): 841–50.

Yang, C H, M a Simon, and H McNeill. 1999. “Mirror Controls Planar Polarity and Equator Formation Through Repression of Fringe Expression and Through Control of Cell Affinities.” Development (Cambridge, England) 126 (24): 5857–66.

Yewale, Chetan, Dipesh Baradia, Imran Vhora, Sushilkumar Patil, and Ambikanandan Misra. 2013. “Epidermal Growth Factor Receptor Targeting in Cancer: A Review of Trends and Strategies.” Biomaterials 34 (34). Elsevier Ltd: 8690–8707. doi:10.1016/j.biomaterials.2013.07.100.

Yosef, Nir, Alex K Shalek, Jellert T Gaublomme, Hulin Jin, Youjin Lee, Amit Awasthi, Chuan Wu, et al. 2013. “Dynamic Regulatory Network Controlling TH17 Cell Differentiation.” Nature 496 (7446): 461–68. doi:10.1038/nature11981.

Yuva-Aydemir, Yeliz, Ann-Christin Bauke, and Christian Klämbt. 2011. “Spinster Controls Dpp Signaling during Glial Migration in the Drosophila Eye.” The Journal of Neuroscience: The Official Journal of the Society for Neuroscience 31 (19): 7005–15. doi:10.1523/JNEUROSCI.0459-11.2011.

Zielke, Norman, Bruce A Edgar, and M. L. DePamphilis. 2013. “Endoreplication.” Cold Spring Harbor Perspectives in Biology 5 (1): a012948–a012948. doi:10.1101/cshperspect.a012948.

Zinzen, Robert P., Charles Girardot, Julien Gagneur, Martina Braun, and Eileen E. M. Furlong. 2009. “Combinatorial Binding Predicts Spatio-Temporal Cis-Regulatory Activity.” Nature 462 (7269): 65–70. doi:10.1038/nature08531.

